# Discretised flux balance analysis for reaction-diffusion simulation of single-cell metabolism

**DOI:** 10.1101/2023.08.01.551453

**Authors:** Yin Hoon Chew, Fabian Spill

**Affiliations:** School of Mathematics, University of Birmingham, Edgbaston, Birmingham, B15 2TT, England, United Kingdom

**Keywords:** flux balance analysis, grid-based, metabolism, reaction-diffusion, spatial method, cell shape

## Abstract

Metabolites have to diffuse within the sub-cellular compartments they occupy to specific locations where enzymes are, so reactions could occur. Conventional flux balance analysis (FBA), a method based on linear programming that is commonly used to model metabolism, implicitly assumes that all enzymatic reactions are not diffusion-limited though that may not always be the case. In this work, we have developed a spatial method that implements FBA on a grid-based system, to enable the exploration of diffusion effects on metabolism. Specifically, the method discretises a living cell into a two-dimensional grid, represents the metabolic reactions in each grid element as well as the diffusion of metabolites to and from neighbouring elements, and simulates the system as a single linear programming problem. We varied the number of rows and columns in the grid to simulate different cell shapes, and the method was able to capture diffusion effects at different shapes. We then used the method to simulate heterogeneous enzyme distribution, which suggested a theoretical effect on variability at the population level. We propose the use of this method, and its future extensions, to explore how spatiotemporal organisation of sub-cellular compartments and the molecules within could affect cell behaviour.

## 1 Introduction

Cellular metabolism is a set of biochemical reactions occurring inside living cells that supports growth and maintenance. Nutrient metabolites are transported into cells, then converted through a series of reactions into energy and building blocks of macromolecular components that make up cellular biomass. Reactions generally occur at regions where enzymes catalysing the reactions are localised. Therefore, metabolites have to diffuse to where enzymes are located to participate in reactions.

In cells, diffusion occurs as molecules collide constantly with one another in a crowded environment. In time, these collisions generate a net movement for each molecular species, from regions where its concentration is high to regions where its concentration is low. Therefore, factors such as the mass and size of the molecular species, and how crowded the environment is, affect the rate of diffusion (Minton, 2001; Gershon et al, 1985; Luby-Phelps et al, 1987). Metabolites are relatively small so they tend to have higher rates of diffusion compared to large molecules like enzymes. However, the effect of metabolite diffusion on the overall rate of reaction depends on diffusion rate relative to enzyme catalytic rate (Schnell and Turner, 2004). Consider the example of a simple single-substrate enzyme-catalysed reaction. Substrate (metabolite) diffusion and catalysis occurs in serial, so the overall rate of reaction is limited by the slower of the two events. When diffusion occurs much faster than enzyme catalysis, the reaction is said to operate at the reaction-limited regime. On the other hand, the reaction is said to operate at the diffusion-limited regime if diffusion is much slower than enzyme catalysis. Which regime any particular reaction operates at may be dynamic because the sub-cellular environment in cells can vary a lot among different cell types and change as the cells grow. Interestingly, several recent studies suggested that some cells may have evolved to achieve optimal growth rates by adopting a molecular composition that provides an optimal (macro)molecular crowding; so as to balance a concentrated environment that supports high catalytic rates and room for movement that enables diffusion (Vazquez, 2010; Oldewurtel et al, 2021; Pang and Lercher, 2023; Dourado et al, 2021). Modelling provides a useful tool to study complex reaction network like metabolism. One established method for modelling metabolic reactions is ordinary differential equations (ODEs). However, this method requires enzyme kinetic parameters whose values are only available for a small fraction of metabolic reactions, such as those in the central carbon metabolism (Teusink et al, 2000; Peskov et al, 2012; Jahan et al, 2016). This limits the use of ODEs for modelling large metabolic network. To circumvent the lack of kinetic measurements, flux balance analysis (FBA) has been commonly used especially for modelling metabolism at the genome scale (Orth et al, 2010). FBA uses linear programming to solve for the rates of reactions by optimising an objective function (that represents an assumed biological objective in each particular study), constrained by measurements such as nutrient uptake, gene expression and/or enzyme concentration. Due to the way linear programming works, reaction rates in the solution may not be unique or obey kinetic rate laws, but analysis of the solution space can provide useful information about the system. While the above are generic methods that can be adopted to model detailed events in metabolism, many metabolic modelling using these methods focus only on the reaction events. Some models consider multiple events and lump them into simple effective functions to represent the overall rate but they also implicitly ignore effects of diffusion. For example, a common rate law used in ODE metabolic models is the Michaelis-Menten equation and its parameters are usually measured *in vitro*, which does not represent the crowded environment in cells where metabolite diffusion is slowed down (Chang et al, 2021; Wittig et al, 2018; Srinivasan, 2022). Genome-scale FBA models, on the other hand, are mostly concerned with (enzyme) capacity that constrains the rate instead of the exact rate because kinetic parameters are not easily available (Blazier and Papin, 2012; Machado and Herrgard, 2014). Estimation from various studies, including analysing the catalytic efficiency of *>*1000 different enzymes, suggested that ignoring diffusion effects may be valid in most cases because enzymatic reactions tend to occur at much lower rates compared to the diffusion of metabolites (Sweetlove and Fernie, 2018; Bar-Even et al, 2015). Thus, it is believed that most metabolic reactions are not diffusion-limited. However, there exists a small number of highly efficient enzymes, where reactions occur very fast such that metabolite diffusion becomes limiting (Pettersson, 1992; Sweetlove and Fernie, 2018). Diffusion-limited effects are exacerbated if the enzymes are in low abundance and heterogeneously distributed within particular cell shapes that require metabolites to travel a long distance or a narrow path (Kinsey et al, 2011; Szymanska et al, 2015).

Various approaches have been used to consider diffusion effects, directly and indirectly, in the modelling of metabolism. These include creating separate rate laws for diffusion-limited and reaction-limited regimes, adjusting the Michaelis-Menten constant to incorporate effects of molecular crowding, and developing a general theory of growth rate optimisation that balances mass conservation, reaction kinetics and limits on cellular density (that can affect diffusion) (Vazquez, 2010; Pang and Lercher, 2023; Dourado and Lercher, 2020). These approaches made theoretical predictions of protein content and cellular density that would be optimal for growth, and it was found that the predicted values fell in the range of experimental measurements in bacterial cells such as *Escherichia coli*. This suggests that the principle of optimality that is assumed in methods such as FBA may be valid, at least under certain growth conditions. Additionally, the approaches above provide an improved representation of metabolism that occur in a crowded environment in cells. They have been useful for studying the quantitative relationship between molecular composition (hence diffusion indirectly) and growth at the cell scale. However, they are not suitable for exploring diffusion effects brought about by spatial properties such as cell shape and heterogeneous distribution of molecular species in the cell. Studies suggested that these effects are not always negligible (Kinsey et al, 2011). A common method for spatial modelling that accounts for both reaction and diffusion is partial differential equations (PDEs). Reaction is usually modelled using the law of mass action that relates reaction rate to substrate concentration, while diffusion is usually modelled using Fick’s first law that relates diffusion rate to concentration gradient. Thus like ODEs, this method requires parameters such as reaction rate constants and diffusion coefficients. Acquiring their values is still a non-trivial problem. A recent work that used PDEs to model the energy metabolism in glial cells proposed a new method to enable simulation of complex cell shapes, and they intentionally left out parameter identification for this very reason (Farina et al, 2021).

To help circumvent the difficulty in obtaining parameter values, we present a method for spatial modelling of reaction-diffusion in single-cell metabolism that has low parameter requirements. Specifically, we expand FBA spatially, add representation of intracellular metabolite diffusion directly to the reaction set, and use linear programming to solve for both the rates of reaction and diffusion that would yield an optimal value for an objective function. Thus, our method embeds diffusion into FBA and only employs linear programming. In this manner, our work differs from existing FBA methods that also extend to implement diffusion, because they coupled FBA with PDE or other methods to represent diffusion outside of the linear programming framework (Henson, 2015; Chen et al, 2016; Borer et al, 2019; Dukovski et al, 2021). Additionally, these methods were developed to consider diffusion in the extracellular environment whereas our method simulates the diffusion of metabolites within an individual living cell. Note that our method is not an alternative to PDE or similar spatial modelling method, but an option for theoretical exploration when rate laws and data such as concentration and kinetic measurements are scarce. In the next section, we provide a description of our method. Then, we describe two example toy metabolic networks and the simulation settings that we used to test and demonstrate the utility of the method; followed by a discussion of the simulation results and their implications. We also suggest potential extension to this method for future work.

## 2 Discretised flux balance analysis

In its basic form, FBA assumes that metabolite concentrations are at steady state, which is generally valid because metabolism operates at a much faster time scale compared to other cellular processes such as transcription and translation (Orth et al, 2010). With this assumption, one can write the mass balance for each metabolite *m* as the following linear equation:

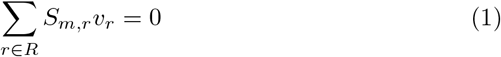

where,

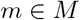

*M* is the set of all metabolites and *R* is the set of all reactions in the model. *S*_*m,r*_ represents the stoichiometric coefficient, which indicates the number of molecules of metabolite *m* that participate in reaction *r* relative to other participants. The stoichiometric coefficient takes a negative sign if the metabolite is a substrate in the reaction, and a positive sign if it is a product. *v*_*r*_ is the flux, i.e. the conversion rate of a substrate to product metabolites, of reaction *r*. Equation (1) can also be represented mathematically in matrix form as follows:

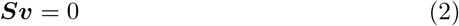

where ***S*** is a matrix of stoichiometric coefficients with each row representing a metabolite and each column representing a reaction (Fig. 1c). ***v*** is then a column vector with each element representing a reaction flux variable. Like ODEs, FBA can be used to solve compartmental models by treating the same metabolite as a distinct species for each of the different compartments (or organelles) that the metabolite occupies; and incorporating transport event between compartments. Therefore, *R* includes not only biochemical reactions, but also metabolite transport; from extracellular compartment into the cell through the plasma membrane, or from one sub-cellular compartment to another. In this case, transport flux variable is the rate of metabolite movement between compartments, akin to “converting” a metabolite in one compartment to the same metabolite in another compartment. Thus, the stoichiometric coefficient takes the value of one, with a positive sign for metabolite in the target compartment and a negative sign for the origin compartment. When modelling single-cell metabolism, flux is in units of particle number (usually moles) per unit of time per cell.

**Fig. 1.**
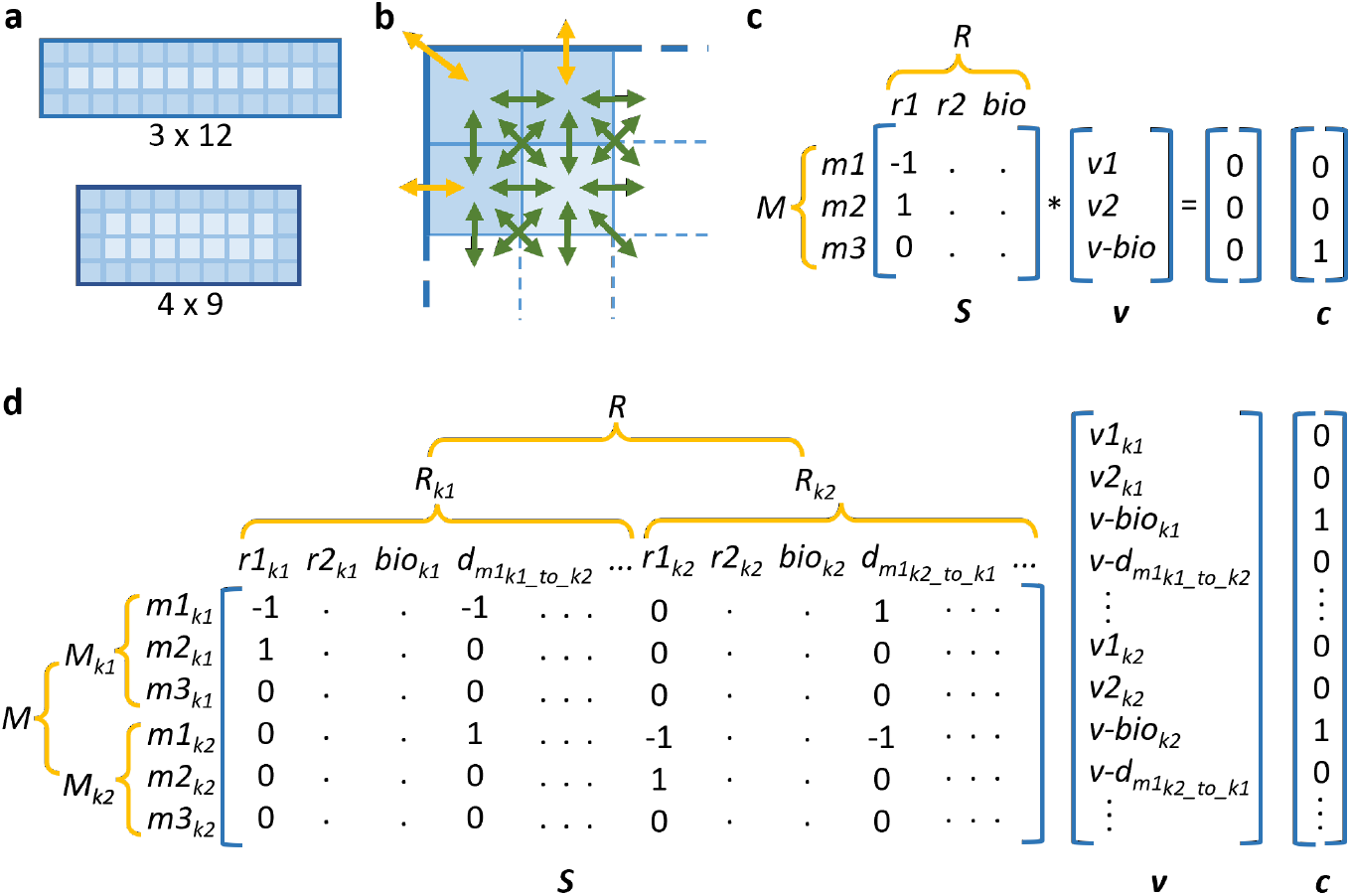
a) In discretised flux balance analysis (FBA), a cell is discretised into a two-dimensional *I* x *J* grid, where *I* represents cell width and *J* represents cell length. Each square element on the grid is a cell region *k*. Cell regions can be grouped into layers, from cell boundary to the centre of the cell; b) Metabolites can diffuse in and out of each cell region to its neighbouring regions (green arrows). For regions at the boundary of a cell, metabolites can also be imported and exported to the extracellular environment (yellow arrows); c) In FBA, the metabolic network is represented as an *M* x *R* matrix of stoichiometric coefficients ***S***, where each row represents a metabolite and each column represents a reaction. ***v*** is a column vector representing reaction flux variables. FBA assumes that the network is at steady state, and solves for the variables by optimising an objective function. ***c*** is a vector of weight for each variable in the objective function. In this example, *v-bio* is the objective function to be optimised; d) Discretised FBA expands conventional FBA by creating regional metabolites *M*_*k*_ and reactions *R*_*k*_ that also include metabolite diffusion *d* between regions. ***v*** and ***c*** are also expanded accordingly.

In this work, we expand FBA to enable spatial modelling within a single cell and include intracellular metabolite diffusion. To spatially represent a cell, we first simplify the representation of cell shape by assuming that each living cell is a rectangle. We discretise a cell into a two-dimensional *I* x *J* grid, where *I* (number of rows) represents cell width and *J* (number of columns) represents cell length (Fig. 1a). We term each square element on the grid a cell region *k*, which has the properties *i* and *j* that denote its coordinate on the grid. Similar to compartmental models, we treat the same metabolite as a distinct species for each of the cell regions. Thus, each cell region contains its own sets of metabolites *M*_*k*_ and reactions *R*_*k*_. Conventional FBA explicitly represents transport of metabolites between compartments by including these events as part of the reaction set. In our discretised cell, intracellular diffusion can be considered a form of passive transport between cell regions. We can therefore add representation of intracellular metabolite diffusion as a natural extension to FBA. We do this in the following manner: 1) for every cell region *k*, movements of metabolites to and from its neighbouring regions are represented as diffusion events and added to *R*_*k*_; 2) if a cell region is located at the boundary between cell and the extracellular compartment, movements of metabolites through the plasma membrane are represented as transport events that are also included in *R*_*k*_ (Fig. 1b). Note that our method considers the extracellular compartment as a single homogeneous region outside of the grid system, therefore only one transport per metabolite is created for each boundary cell region. Similar to a transport event, the stoichiometric coefficient for a diffusion event takes the absolute value of one, and diffusion flux variable is the rate of net metabolite movement between cell regions. The new sets of metabolites *M* and reactions *R* for the whole cell are now the union of all cell-region sets *M*_*k*_ and *R*_*k*_, respectively. Our method therefore expands the number of elements in the matrix of the original FBA problem, by decomposing each metabolite and reaction flux at the whole-cell scale into their constituents at the cell-region scale, and incorporating diffusion fluxes between regions (Fig. 1d). Flux in this case is in units of particle number (or moles) per unit of time per cell region.

Metabolic network usually contains a higher number of reactions than the number of metabolites. This creates an under-determined system where there are more unknown fluxes than there are equations. FBA uses linear programming to determine the unknown fluxes by adding inequality constraints and optimising an objective function. The linear programming problem can be formulated as follows:

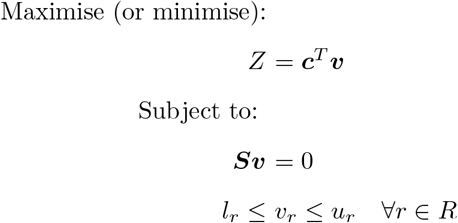

Here, ***c*** is a vector of weights, or objective function coefficients, that is used to indicate the level of contribution of each flux to the objective function *Z*. A common objective function in modelling genome-scale metabolic network is the biomass reaction, which is a pseudo-reaction that represents cell growth (see example in Section 3.1). This can be indicated by setting the weight of biomass reaction to one and all others to zero. As our method decomposes cell-scale flux variables into regional-scale variables, we need to re-compose the latter to generate the equivalent objective function. Using biomass reaction as an example, our method will set the weight of biomass reaction in every cell region to one, so that their fluxes will be added back to give whole-cell biomass growth (Figs. 1c and 1d). The optimised value of the objective function depends on the inequality constraints. Each flux variable *v*_*r*_ is constrained between a lower bound *l*_*r*_ and an upper bound *u*_*r*_. Their values can be set based on experimental measurements or knowledge about the variable. For example, in the case of irreversible reactions that can only occur in the forward direction, one can set their lower bounds to zero. If no data or knowledge is available, variables can be left unbounded, i.e. the lower and upper bounds are set to negative infinity and positive infinity, respectively. For diffusion flux variables, their lower bounds are automatically set to zero because our method creates a separate variable for each direction between any two neighbouring cell regions. Note that FBA does not have any notion of metabolite concentration; flux values in the optimal solution depend on the constraints set in each particular simulation. These constraints eliminate infeasible states to help narrow down the solution space. Within the feasible region, it is possible to have more than one set of solution that yield the same optimal value for the objective function. Depending on the accuracy of the constraints, some of these solutions may not be biologically feasible. Nevertheless, in the absence of parameters such as rate constants and diffusion coefficients, setting the objective function and constraints for flux variables in different ways allows one to theoretically explore the solution space and gain insights into the system (Loghmani et al, 2022).

Note that in general, FBA does not have explicit representation of mass, volume and surface area. The scale of the system to be modelled is indicated by the units of the constraints. Therefore, changing the number of cell regions, i.e. grid elements, does not necessarily indicate a change in cell size in our method. For example, one can increase the number of grid elements but sets the units of the constraints such that the sum over all regions will recover the same cell-scale units and values. Doing this does not change cell size but increases the spatial resolution of the simulation. One advantage of having spatial resolution is that we can use different constraints for the same reaction in different regions. This can be useful for studying local effects and how they influence the whole system.

Our method has been implemented as a Python API. Codes for the method, as well as the following toy models and simulations, are available on GitHub at https://github.com/YinHoon/discretised FBA. Our API uses COBRApy, a constraint-based modeling package that is commonly used for running conventional FBA (Ebrahim et al, 2013).

## 3 Model description

In general, genome-scale metabolic modelling using FBA requires two elements: 1) a metabolic network; 2) a set of constraints and objective function that turns the network into a simulable model. Metabolic networks are composed of interconnected pathways; and a pathway in its simplest form can be described as the path, made of a series of biochemical reactions, that a starting metabolite takes and ends with a final product. The metabolic network of an organism can be reconstructed from its genomic information. By comparing to a reference genome, we can infer which established enzyme-catalysed reactions likely take place in the organism based on the existence of genes that express the associated enzymes. Typically, individual steps within enzyme-catalysed reactions, such as the formation of enzyme-substrate complex and the release of product, are not part of metabolic networks. Instead, genes (and indirectly the enzymes that they express) are only incorporated as metadata associated to the relevant reactions. These metadata may be used when determining the constraints of flux variables. For example, if gene expression data indicate that some genes have very little expression, the flux variables for reactions associated to these genes can be constrained to smaller values to reflect reduced enzyme capacities. In terms of setting up an objective function, the underlying assumption in FBA is that cells have evolved through selective pressure to achieve an optimal rate of specific cellular functions. For example, bacteria and cancer cells are assumed to operate at metabolic states that maximise their growth rate, while muscle cells may be maximising energy production. The validity of these assumptions is open to debate, but they have provided solutions that best matched measured data in various contexts (Costa et al, 2014; Schuetz et al, 2007; Schnitzer et al, 2022).

To test our method, we created two toy models that exemplify two common types of metabolic pathways. The first is a linear pathway, where a metabolite is converted into one final product through a series of intermediate forms. The second is a branched pathway, where the path diverges and an intermediate can go on to form different final products.

### 3.1 Example 1: A linear pathway

In this section, we first present the model at the cell scale, i.e. the set of reactions (metabolic network) and parameters (constraints and objective function) at the organism level that is normally represented when using conventional FBA. We then describe how our method transforms cell-scale representations into regional-scale representations that include region-specific reactions and metabolite diffusion.

The toy metabolic network consists of only a few reactions, as follows:

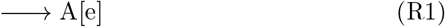

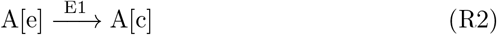

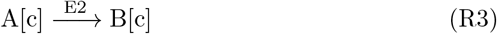

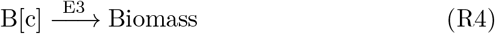

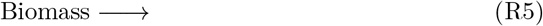

The network of reactions occurs in two compartments (as denoted in square brackets): an extracellular compartment “e” and a cellular compartment “c”. In reaction R2, metabolite A is transported from the extracellular compartment into the cellular compartment via a transporter E1. Metabolite A is then converted into B in the cell, catalysed by enzyme E2. Finally, B is converted into cell biomass, catalysed by enzyme E3. R1 and R5 are exchange reactions, i.e. pseudo-reactions that act as the source and sink to balance out all other reactions in the network. R5 is the biomass reaction, a proxy of cell growth, that will be set as the objective function and maximised during simulation. All reactions are irreversible.

Given the above reaction network, conventional FBA converts it into the following matrix-vector equation of the product of ***S*** and ***v*** to represent a system at steady state:

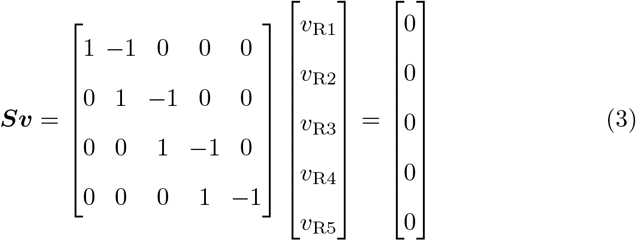

The rows in ***S*** represent the four metabolite species in the network: “A[e]”, “A[c]”, “B[c]” and “Biomass”. For example, one molecule of metabolite species “A[e]” is produced through reaction R1 but consumed through reaction R2. This is represented in the first row of ***S***, where the values of 1 and -1 are entered in the first two columns. Since “A[e]” does not participate in the rest of the reactions, entries for the rest of the columns in this row are zeroes. The objective function is to maximise *v*_R5_, which can be represented in vector form as follows:

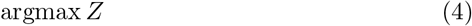

where

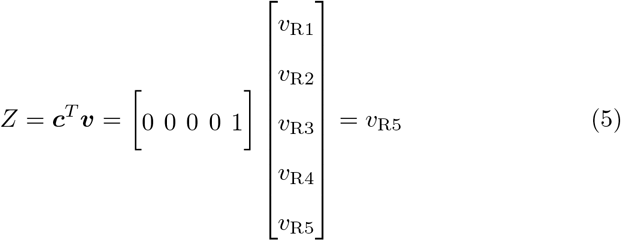

Finally, inequality constraints are added to complete the linear programming problem. These constraints determine the range of values that each flux variable is allowed to take, and the solver will eliminate sets of solution that violate the constraints even if they satisfy equations (3) – (5). Common types of data for constraining conventional FBA models include nutrient uptake rates, fluxomic, transcriptomic and proteomic data (Blazier and Papin, 2012; Machado and Herrgard, 2014; Cook and Nielsen, 2017). When using transcriptomic and proteomic data, one assumption made is that the level of gene expression indicates maximum enzyme capacity which limits the rate of reaction. For our simulation, we chose to constrain the fluxes of reactions R2, R3 and R4 whose rates are limited by the amount and efficiency of E1, E2 and E3, respectively. We picked these reactions so that we could demonstrate the utility of spatial resolution in our method (see next section for details). Since the reactions are irreversible, we constrained the reaction fluxes by setting their lower bounds to zero while their upper bounds are determined based on maximum enzyme capacity. This can be written as the following inequality:

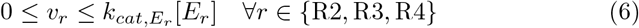

where *k*_*cat*_ is the catalytic constant, in per unit of time, and [*E*_*r*_] is the concentration of enzyme (transporter) *E* that catalyses (facilitates) reaction *r*. We employed this upper-bound expression because it is commonly used in enzyme kinetic studies to represent maximum enzyme capacity (Davidi et al, 2016; Heckmann et al, 2018, 2020). Here, enzyme concentration is expressed as particle number per cell, i.e. total cellular amount, in accordance with the units of fluxes. We set the total cellular amount of each enzyme (or transporter) to 100 and their *k*_*cat*_ values to 1.0. We further set the upper bound for the flux of exchange reaction R1 to 100, as a proxy of maximum rate of nutrient provision, as follows:

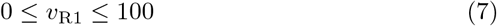

For the flux of biomass reaction R5, its upper value was left unbounded and to be maximised in the objective function, while its lower bound was set to zero to ensure a positive growth, as follows:

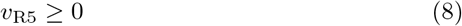

We chose the above set of parameter values so that the maximum feasible growth per cell is 100 and no specific enzyme is limiting growth. Note that we are not assigning specific units in this toy example, but the parameter values are to be taken as being consistent in units.

Given a cell-scale representation such as the one described above, our method converts it into a regional-scale representation using the following steps (see Appendix A for more details on using the Python API):

1. Create a two-dimensional *I* x *J* grid containing a set of regions, *K*, that define the spaces occupied by the individual living cell;
2. Expand the set of metabolites by decomposing each metabolite species in the cellular compartment into regional species, and create the metabolite objects for every cell region. For example, an object “A[c]_*k*_” is created for each cell region *k* to represent metabolite A in that region. The same is done for metabolite B and biomass. Doing this increases the number of rows in the matrix ***S*** in equation (3). For metabolite A that also exists in the extracellular compartment “e”, only one object “A[e]” is created since this compartment is treated as a single region outside of the grid system;
3. Expand the set of reactions by decomposing each biochemical reaction that occurs in the cellular compartment into regional reactions, and create the reaction objects for every cell region. For example, an object “R3_*k*_” is created for each cell region *k* to represent reaction R3 in that region. This reaction converts one molecule of metabolite “A[c]_*k*_” to one molecule of “B[c]_*k*_”. Thus, the stoichiometric coefficients 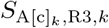 and 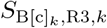 are assigned the values of -1 and 1, respectively. A variable *v*_R3,*k*_ is generated to represent the flux through this reaction event. This increases the number of columns in the matrix ***S*** and the size of vector ***v*** in equation (3). Since reaction R1 only involves metabolites in the extracellular compartment, only one reaction object “R1” is created;
4. Create an objective function that is mathematically equivalent to the cell-scale objective function, by recomposing over all cell regions. In the toy model presented above, the objective is to maximise the (total cellular) flux through reaction R5. The recomposed objective function is therefore:

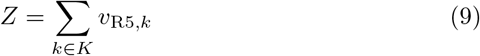
5. Further expand the set of reactions (and the size of ***S*** and ***v***) by decomposing each cross-membrane transport process into regional transport processes, and create the reaction objects for every cell region located at the boundary between cell (grid system) and the extracellular compartment (outside the grid system). For example, an object “R2_*k*_” is created for each boundary cell region *k* to represent the transport process R2 between that region and the extracellular compartment. This reaction “converts” one molecule of metabolite “A[e]” to one molecule of “A[c]_*k*_”. As in previous, the stoichiometric coefficients *S*_A[e],R2,*k*_ and 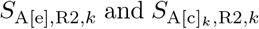 are assigned the values of -1 and 1, respectively, and a variable *v*_R2,*k*_ is generated to represent the flux through this transport event;
6. Further expand the set of reactions (and the size of ***S*** and ***v***) by creating new reaction objects that represent the diffusion process of each metabolite species between each pair of neighbouring cell regions. For example, to represent the diffusion of metabolite A from cell region *k*_1_ to a neighbouring region *k*_2_, a reaction object 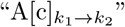 is created. Just like a transport event, the stoichiometric coefficients 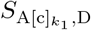 (where D is a shorthand for the diffusion identifier) and 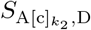 are assigned the values of -1 and 1, respectively, and a variable *v*_D_ is generated to represent the flux through this diffusion event;
7. Set the bounds that constrain the flux variables generated in steps 3, 5 and 6. For reactions R2, R3 and R4, [*E*_*r*_] in inequality (6) represents the total cellular amount that limits the total cellular flux of reaction *r*. Since each cell region *k* only contains a portion of the total cellular enzyme, the regional flux is capped by the regional portion [*E*_*r*_]_*k*_. This can be expressed as:

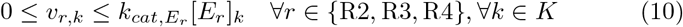

For reaction R5, the lower bound for each regional flux is set to zero to ensure a positive growth everywhere:

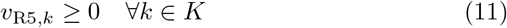

For reaction R1 that has not been decomposed, its flux variable is bounded as in inequality (7). For the diffusion flux variables generated in step 6, we used different bounds to simulate different rate-limiting regimes (see details in Section 4).

### 3.2 Example 2: A branched pathway

Branched pathways are basically (linear) pathways that intersect at a branch point. They are prevalent in metabolic networks and methods continue to be developed to identify them (Huang et al, 2021). One prominent branch point in metabolism is pyruvate, which is the end-product of the linear glycolysis pathway that starts from glucose. Pyruvate can move on to produce energy efficiently through oxidative phosphorylation, or it can be converted to lactate and produce energy quickly but less efficiently under certain conditions, such as during low oxygen availability. Pyruvate and lactate metabolism have great implications on human health; they have been associated with conditions such as cancer, inflammation, neurodegeneration and heart failure (Li et al, 2022; Gray et al, 2014). In this example, we created a simple branch pathway that encapsulates some of the scenarios above, as follows:

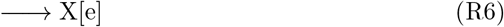

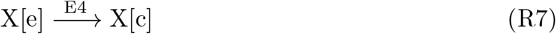

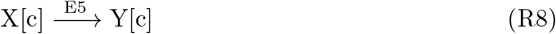

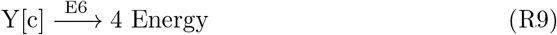

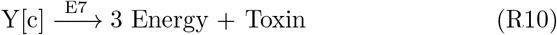

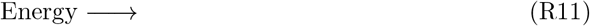

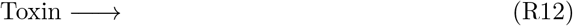

As in the previous example, this network of irreversible reactions also occurs in two compartments. Metabolite X is first transported from the extracellular compartment into the cellular compartment via a transporter E4. Metabolite X is then converted into Y in the cell, catalysed by enzyme E5. From this point, metabolite Y can participate in reaction R9 or R10. In reaction R9, each molecule Y can produce four units of energy through catalysis by enzyme E6. In reaction R10 that is catalysed by enzyme E7, each molecule Y can produce three units of energy and one molecule of a compound that can be toxic to the cell. Again, pseudo-reactions R6, R11 and R12 were added to balance out all other reactions in the network. For this model, we would be combining two objectives: 1) maximising energy production; 2) minimising toxin production. Again, we set the total cellular amount of each enzyme (or transporter) to 100. The *k*_*cat*_ values for all enzymes were set to 1.0, except enzyme E7 which has a *k*_*cat*_ of 2.0. We further set the upper bound of exchange reaction R6 to 100, to represent maximum rate of nutrient provision.

With this reaction network, the steady-state equation and inequality constraints for a cell-scale representation become:

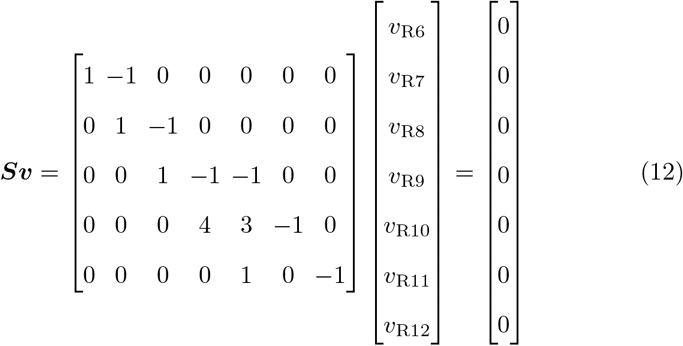

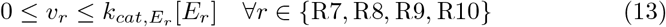

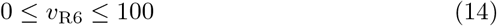

To ensure that reactions R11 and R12 have positive fluxes, the following inequality is used:

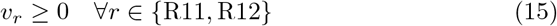

Since this model has two objectives, we would need to perform multi-objective optimisation and there are various ways this can be done (Gunantara, 2018). Here, we chose to use linear scalarisation, where multiple objectives are linearly combined into a single objective function and a weight is assigned to each objective term. In the current case where flux variable *v*_R11_ is to be maximised while *v*_R12_ is to be minimised, we used positive and negative weights, as follows:

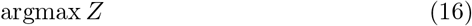

where

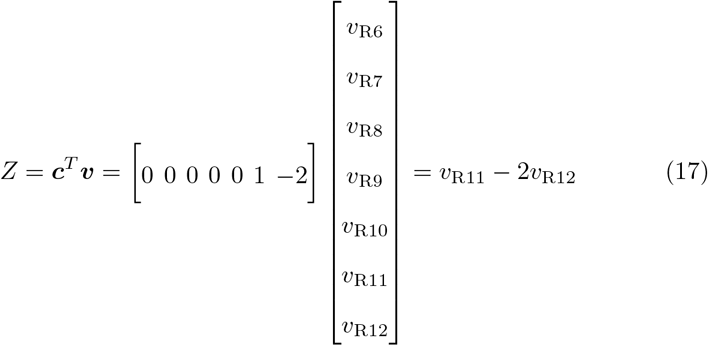

We assigned a higher absolute value to the weight of *v*_R12_ to exert a strong penalty on toxin production. With this value, it is generally more favourable to direct all molecules Y to be converted to energy through reaction R9.

Using the same steps as described for the previous example, a regional-scale representation can be created by decomposing flux variables into regional variables, expanding ***S*** and ***v*** accordingly, adding new flux variables for diffusion, constraining regional variables with local limits (e.g. based on regional portions of the total cellular enzyme; see next section for details), and recomposing the flux variables in the objective function.

## 4 Simulation settings

We conducted a series of simulations using the two examples described above, with the aim of testing our method and its utility. First, we wanted to check that our method has the ability to reproduce the expected effects of diffusion. Even though we have expanded conventional FBA to explicitly represent events of intracellular metabolite diffusion, their fluxes are solved using linear programming that depends on the objective function and set of constraints used. There is no guarantee that the solution(s) will capture the diffusion phenomenon at all. To test this, we ran simulations of different cell shapes, with intracellular reactions operating at either reaction-limited or diffusion-limited regime. To simulate a reaction-limited regime (*v*_*r*_ *≪ v*_D_), we left the diffusion flux of substrate in the reaction unbounded, i.e. the substrate diffuses instantaneously. To simulate a diffusion-limited regime, (*v*_*r*_ *≫ v*_D_), i.e. substrate diffusion is negligible, there are two ways this can be done. One can constrain every flux variable *v*_D_ to be equal zero, or one can omit diffusion from the set of reaction objects and flux variables all together. We chose to do the latter since it reduces the size of the linear programming problem.

Second, we wanted to test the utility of the method for simulating spatial heterogeneity within a single cell. It is known that the sub-cellular environment is heterogeneous, which means that enzymes are likely not distributed uniformly. However, other than the confinement of specific enzymes within certain organelles, it is unclear if there are distinct enzyme distribution patterns, how cellular behaviour is affected by these patterns if any, and under what conditions. Our method is well-suited to explore this theoretically since the flux variable for each intracellular reaction is decomposed into regional flux variables, and they can be separately constrained with their own bounds as shown in inequality (10). Using this inequality, we can incorporate the spatial information of enzyme distribution by assigning specific portions of total cellular enzyme to specific regions based on the regions’ spatial location within the cell. This means that we can simulate the local rates of each reaction, as determined by local availability of enzymes, and evaluate the global rate at the cellular level.

To facilitate the exploration of enzyme distribution effect, we have included an additional function in the Python API that allocates enzymes based on user-specified parameters. Currently, we have implemented four options for allocating enzymes in the cell: 1) uniformly; 2) on a positive gradient (increasing concentration from the boundary to the centre of the cell); 3) on a negative gradient or; 4) randomly. The function uses the following to determine the portion [*E*]_*k*_ that each cell region will receive out of the total cellular amount

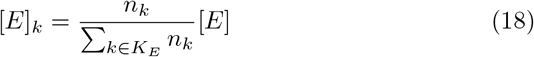

Here, *K*_*E*_ is the set of cell regions where enzymes will be allocated to and *n*_*k*_ is a non-negative value assigned to cell region *k*. If a user chooses to distribute an enzyme uniformly, *n*_*k*_ will be set to one for all cell regions. For positive (negative) gradient distribution, *n*_*k*_ for cell regions in the outermost (innermost) layer of the cell will be assigned a value of one and increase by a user-defined gradient parameter value for each layer into (away from) the cell centre (Fig. 1a). Finally, users can also choose to distribute an enzyme randomly without any specific pattern. In this case, *n*_*k*_ for each cell region is sampled from a random number generator. Once the regional portions of each enzyme have been allocated using equation (18), they can be used to constrain the corresponding flux variables (see Appendix B for more details on using the API).

Third, we wanted to test using the method for spatial stochastic simulation. As mentioned earlier, our method allows us to incorporate the spatial information of enzyme distribution. One type of enzyme distribution we have implemented is random distribution. With a random distribution, the portion of total cellular enzyme that each region gets does not depend on its spatial location within the cell. Instead, each time we simulate a cell with the same shape, the same region *k* will be allocated a different portion because a new value will be sampled for *n*_*k*_ in equation (18). Consequently, the region’s flux variables will be constrained by different values each time. Therefore, compared to the other three distribution patterns that will always yield the same set of constraints and the same solution, a random enzyme distribution could produce a different solution to the linear programming problem in each run. This is akin to simulating a different cell each time we run the simulation. We used this feature to do spatial stochastic simulation.

Altogether, we conducted the three tests above through different combinations of the following parameters:

### Cell shape

1x36, 2x18, 3x12, 4x9, 6x6 (note that the total number of cell regions is the same for every cell shape so they all have the same spatial resolution)

### Rate-limiting regime

no substrate diffusion (all intracellular reactions are diffusion-limited), one substrate diffuses instantaneously while the others cannot diffuse (a mixed of reaction-limited and diffusion-limited reactions), all substrates diffuse instantaneously (all intracellular reactions are reaction-limited)

### Distribution pattern for transporters

uniform, random (we distributed transporters only among cell regions in the outermost layer of the cell since transporters reside on the membrane)

### Distribution pattern for enzymes

uniform, positive gradient, negative gradient, random (we distributed enzymes among all cell regions)

For positive and negative gradients, we set the gradient parameter to 1 and -1, respectively. For stochastic simulations, i.e. ones that involved random enzyme distribution, we ran batches of 100 simulations to determine population-level behaviour. Note that we did not seed the random number generator for these simulations, but we have repeated the batches of simulations multiple times, and our results were robust and showed consistent population-level behaviour. Scripts for running these simulations can be found on the method’s GitHub page. All were steady-state simulations, an inherent feature of FBA. Simulation results are presented next.

## 5 Simulation results

### 5.1 Method captures the effects of diffusion

We first tested whether our method can reproduce the effects of intracellular metabolite diffusion, by comparing simulations across different cell shapes. For this test, we looked at simulations of the linear-pathway example because it has a straightforward single path for mass flux. We also focused only on deterministic simulations, i.e. parameter combinations that do not involve any random enzyme distribution. To allow a quantitative comparison across cell shapes, we used the perimeter-to-area (PA) ratio to represent each shape. We defined this ratio as the number of cell regions in the outermost layer divided by the total number of cell regions. This ratio was chosen over a commonly used aspect ratio because it is the two-dimensional equivalent of surface-area-to-volume ratio, which is known to correlate positively with the rate of diffusion (Okie, 2013). If the effects of diffusion are captured by the method, we would expect to see differences in optimised cell growth at different PA ratios when diffusion becomes limiting. Note that in our simulations, transporter E1 was distributed only to regions in the outermost layer regardless of cell shape. We can therefore rule out cellular capacity for nutrient uptake as the cause of any differences in simulated growth if observed. As seen in Fig. 2, when either or both metabolites A and B have limited ability to diffuse, there is a general trend of reduced growth at low PA ratios. On the other hand, in the control where both metabolites can diffuse instantaneously, i.e. all intracellular reactions are reaction-limited as implicitly assumed in conventional FBA, cell growth can achieve the maximal value of 100 regardless of the ratio. We observed these relations in all combinations of non-random enzyme distributions. It is worth noting that both 1x36 and 2x18 cells achieve the same growth in our simulations despite having different elongated shapes. This is because at this spatial resolution, they have the same PA ratio where all cell regions are in the outermost layer. Each cell region can receive nutrients from the extracellular compartment via transporters instead of being dependent on diffusion from neighbouring regions. A higher spatial resolution would be needed to discriminate between the two shapes, but we included both in our simulations as controls to check that the reduced growth effects we observed in other shapes were indeed due to intracellular diffusion. These results demonstrated that our method can spatially simulate reaction-diffusion in single-cell metabolism. It can capture the theoretical effects of both reaction-limited and diffusion-limited regimes on growth, as well as the spatial effect of cell shape.

**Fig. 2.**
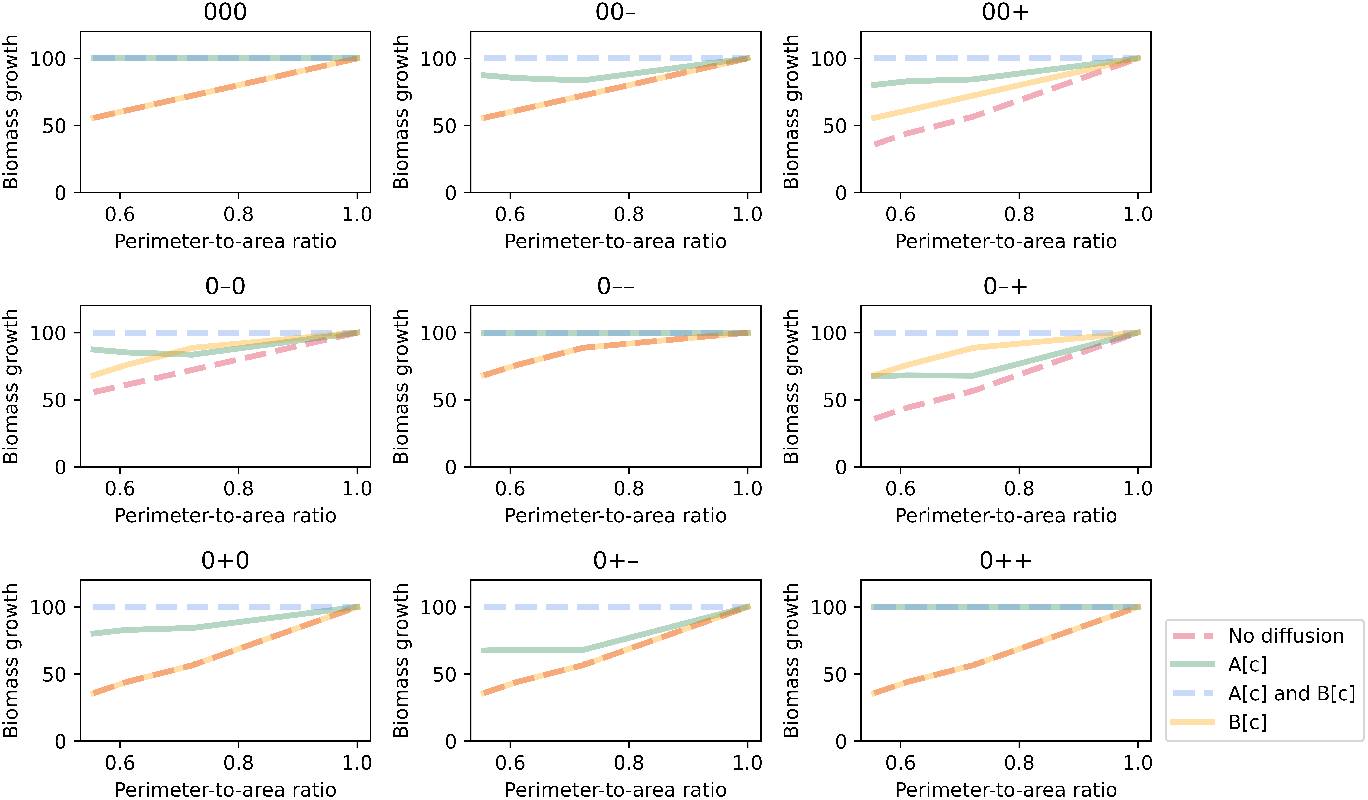
Simulated biomass growth plotted against perimeter-to-area ratio for different rate-limiting regimes (no diffusion, metabolite A can diffuse instantaneously, metabolite B can diffuse instantaneously, both metabolites can diffuse instantaneously). Each panel illustrates plots for a different combination of enzyme distribution patterns, as indicated by three characters at the top of the panel (“0”: uniform distribution; “+”: positive gradient; “-”: negative gradient) that represent the distribution of transporter E1, enzyme E2 and enzyme E3, respectively. Opacity of the lines was adjusted so that overlapping lines appear as a blend of the overlapping colours.

The extent to which optimal growth is reduced with PA ratio, i.e. the slope of lines in Fig. 2, depends on the metabolite(s) that is not diffusible. In general, the effect of cell shape is strongest when both metabolites A and B cannot diffuse. This is because all intracellular reactions are diffusion-limited, so a reduction in diffusion in rounder cells increases the severity of the limitation. The effect is weaker if metabolite A can diffuse instantaneously, since only one reaction is diffusion-limited. However, combination of enzyme distribution also plays a role. We explored this next.

### 5.2 Method enables study of enzyme distribution effect

To inspect how enzyme distribution influences the effect of diffusion on growth, we further analysed simulated growth of a 6x6 cell where diffusion-limiting effect is most severe. Again, we only focused on deterministic simulations where transporter E1 has a uniform distribution, while enzymes E2 and E3 can be distributed uniformly or on a (positive/negative) gradient. We found that for each rate-limiting regime, combinations of enzyme distribution can be grouped based on their effective maximal growth rates (Fig. 3).

**Fig. 3.**
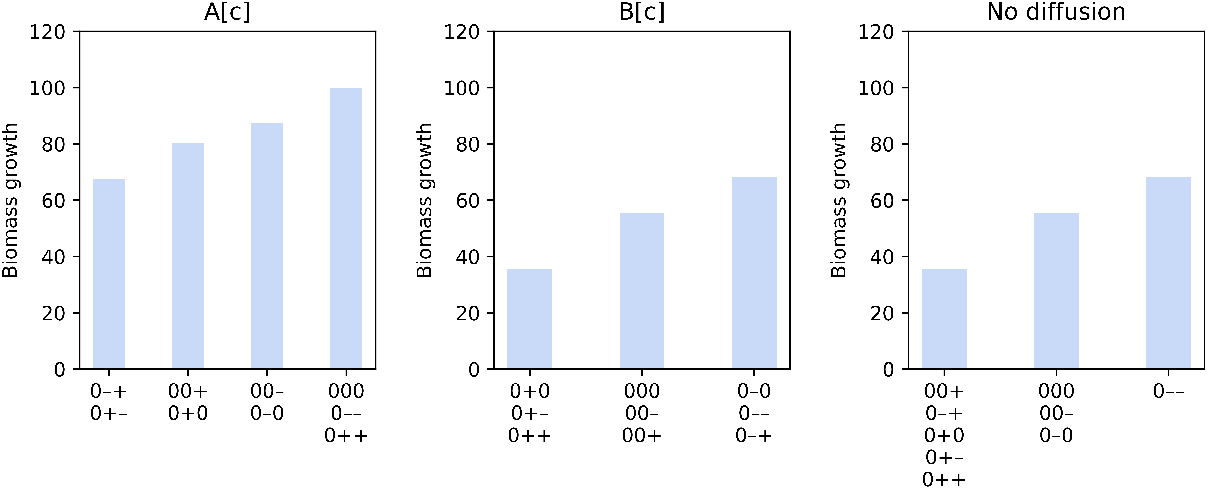
Simulated biomass growth for a 6x6 cell for different rate-limiting regimes: only metabolite A can diffuse instantaneously (left panel); only metabolite B can diffuse instantaneously (middle panel); and both metabolites are restricted (right panel). The combinations of enzyme distribution patterns that produce each growth level are indicated at the bottom of each bar. Each combination of three characters (“0”: uniform distribution; “+”: positive gradient; “-”: negative gradient) represents the distribution of transporter E1, enzyme E2 and enzyme E3, respectively.

For the regime where only metabolite A can diffuse instantaneously (left panel in Fig. 3), a maximal growth of 100 can still be achieved as long as enzymes E2 and E3 have the same distribution pattern, i.e. both enzymes have the same concentration in every cell region. This is because A can move freely and instantaneously so it is optimally channelled in the simulation to occupy each cell region at a rate proportional to regional concentration of enzyme E2. This enables enzyme E2 in every region to operate at maximal capacity, without being limited by diffusion, to convert A to B. If in addition there is proportional amount of regional enzyme E3 to convert B to biomass on site, growth is not limited by the diffusion of B either so the maximal growth is achieved. On the contrary, if the enzymes are distributed in opposing gradient to each other, the growth in every cell region is limited by either of the enzymes. Even if A can diffuse accordingly to enable maximal conversion to B in every region, some regions do not have sufficient E3 to fully support on-site conversion of B to biomass, and any excess B cannot diffuse to other regions with higher E3 to make up for it. Thus, overall growth is lowest for combinations with opposing gradients. In the middle range are cases where one enzyme is uniformly distributed while the other has a gradient distribution. In these cases, a negative gradient is more favourable for growth. This can be explained by the fact that the number of cell regions in each layer decreases from the outer layers to the inner layers (Fig. 1a). With a negative gradient where concentration of an enzyme is higher in the outer layers, more cell regions are allocated with high amount of the enzyme thereby generating higher overall growth. Again, this highlights how PA ratio, metabolite diffusion and enzyme distribution interact to give rise to growth behaviour.

Our analysis also shows that there are only three growth levels when metabolite A cannot diffuse, regardless of whether B can diffuse (middle and right panels in Fig. 3). However, combinations of enzyme distribution are grouped differently depending on the ability of B to diffuse. If B can instantaneously diffuse, combinations are grouped based only on the distribution of enzyme E2. A negative gradient is most favourable, followed by a uniform distribution and lastly a positive gradient. The reason is quite simple. Since metabolite A cannot diffuse, they stay in cell regions in the outermost layer after being transported into cell. A negative gradient means there is a higher concentration of enzyme E2 in the outermost layer to maximise conversion of A to B. Hence, there is more B that can be converted to biomass either immediately in this layer or after diffusing to the inner layers, to produce a higher growth rate. If both metabolites A and B cannot diffuse, reactions and growth only occur in cell regions in the outermost layer. Thus, a high growth rate can only be achieved when both enzymes E2 and E3 have high concentrations in this layer. In accordance with this, growth is lowest when any enzyme has a positive gradient distribution, i.e. a low concentration in the outermost layer.

In the optimal solutions above, for cases with reaction-limited regime, metabolites diffuse in the direction and rate that maximises growth. Each metabolite tends to move towards regions with high enzyme capacity, where it will be consumed at a higher rate and stays at a lower level. If we assume that volume exclusion by other molecules is the same in each region, then the optimal solutions have simulated metabolite diffusion in the direction from regions with high concentrations to regions with low concentrations, without explicitly representing concentration gradient. Note however that the reaction network in this simple example only consists of single-substrate linear pathway. For more complex reaction network, the optimal solution may not necessarily favour diffusion that follows the concentration gradient. Additional constraints may be needed to obtain more accurate solutions.

### 5.3 Effect on cell-to-cell variability

In this section, we focus on simulations that involve random enzyme distribution. Fig. 4 shows the growth distribution of 100 stochastic simulations for each rate-limiting regime and combination of enzyme distribution for a 6x6 cell. As previously observed, the maximal growth of 100 is always achieved when metabolites A and B can diffuse instantaneously. For some rate-limiting regimes, cells can switch between deterministic and stochastic growth, depending on the combination of enzyme distribution. Interestingly, several growth distributions have lower standard deviation than others. To explore this further, we plotted the coefficient of variation (CV) against the PA ratio for each case (Fig. 5). We used CV instead of standard deviation because the former is normalised to the population mean so it is comparable across cases.

**Fig. 4.**
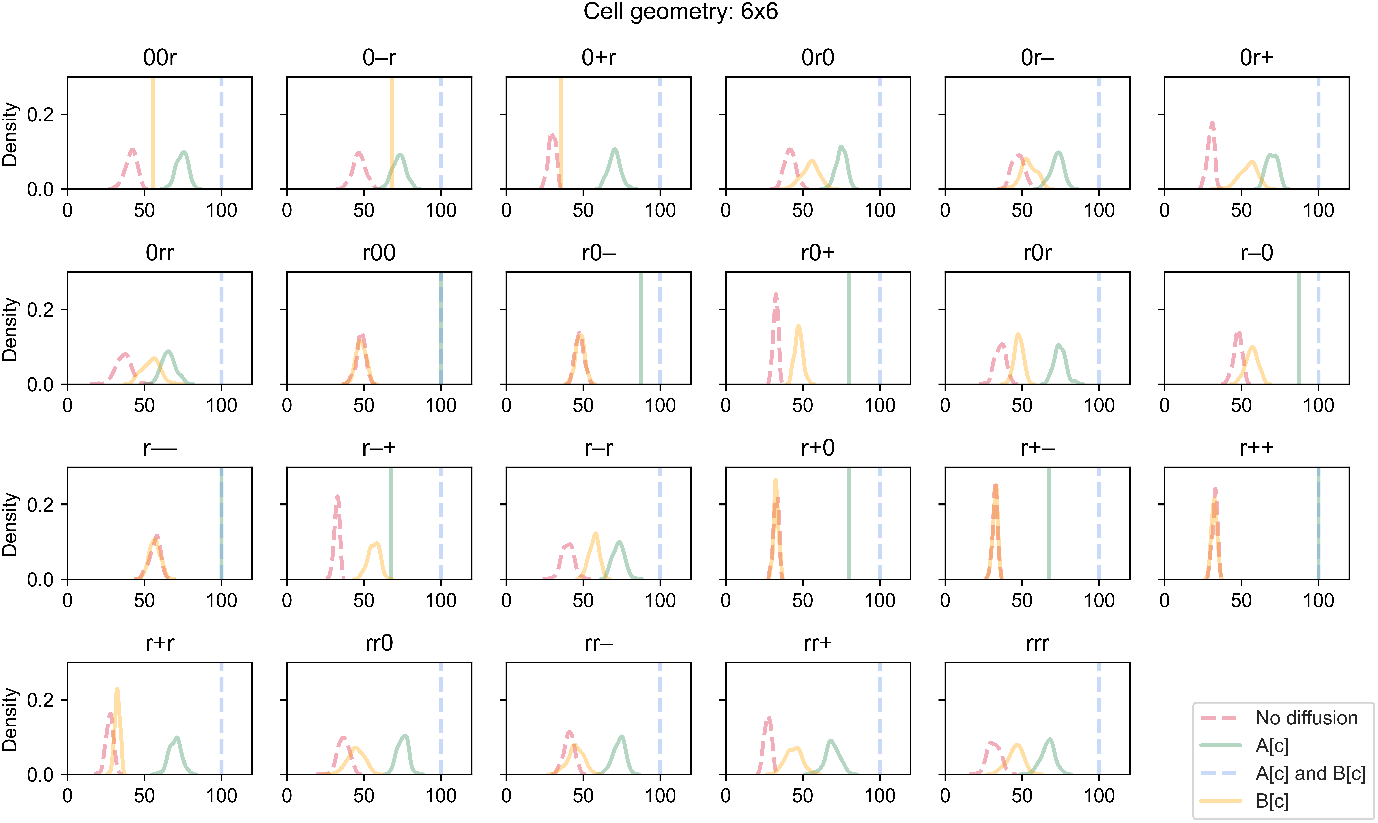
Kernel density plots of simulated biomass growth for different rate-limiting regimes (no diffusion, metabolite A can diffuse instantaneously, metabolite B can diffuse instantaneously, both metabolites can diffuse instantaneously). Each panel illustrates plots for a different combination of enzyme distribution patterns, as indicated by three characters at the top of the panel (“r”: random distribution; “0”: uniform distribution; “+”: positive gradient; “-”: negative gradient) that represent the distribution of transporter E1, enzyme E2 and enzyme E3, respectively. Each plot describes the population distribution of 100 simulations. A vertical line indicates that all 100 simulations produce the same value. Opacity of the lines was adjusted so that overlapping lines appear as a blend of the overlapping colours.

**Fig. 5.**
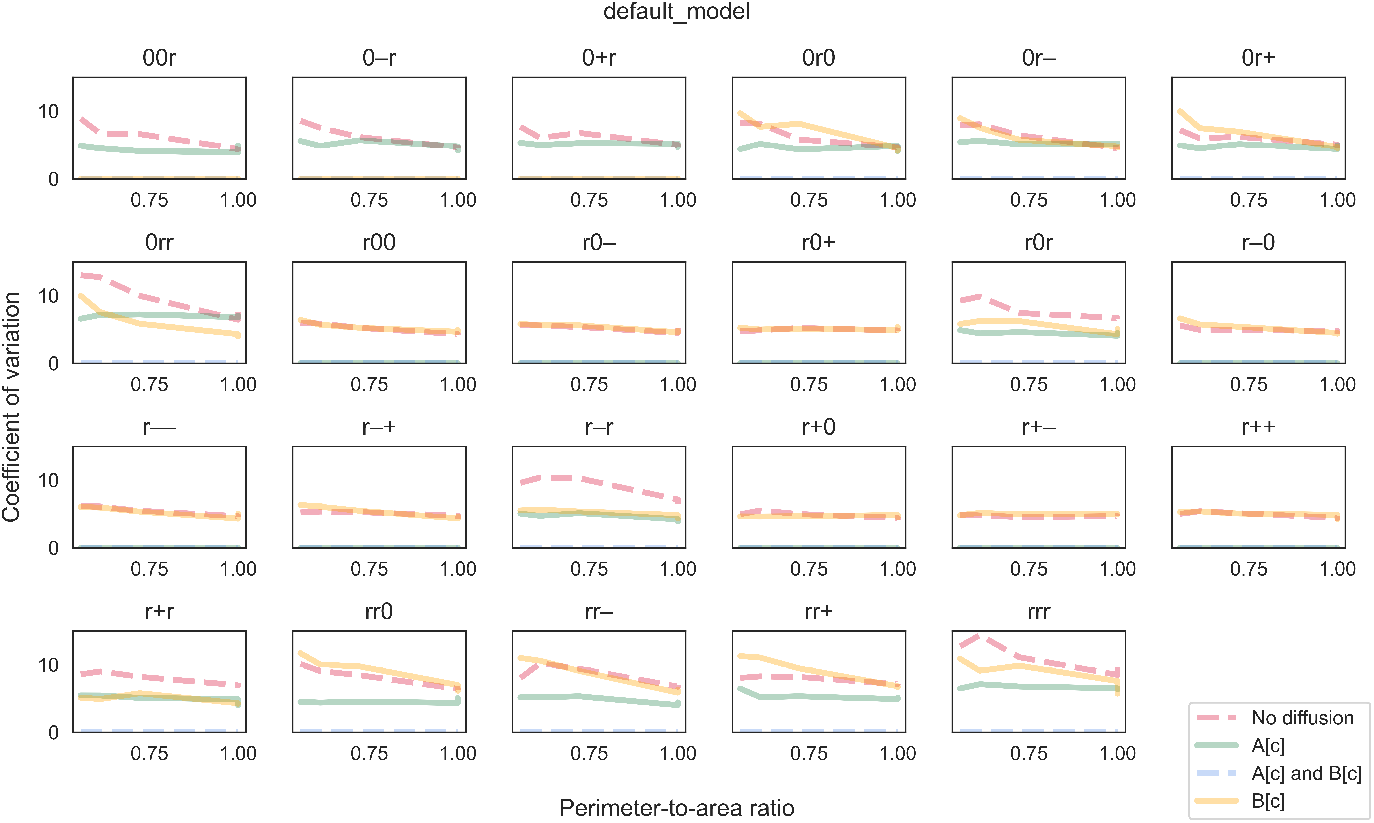
The coefficient of variation (CV) of biomass growth simulated using the “default model”. CV is plotted against perimeter-to-area ratio for different rate-limiting regimes (no diffusion, metabolite A can diffuse instantaneously, metabolite B can diffuse instantaneously, both metabolites can diffuse instantaneously). Each panel illustrates plots for a different combination of enzyme distribution patterns, as indicated by three characters at the top of the panel (“r”: random distribution; “0”: uniform distribution; “+”: positive gradient; “-”: negative gradient) that represent the distribution of transporter E1, enzyme E2 and enzyme E3, respectively. Opacity of the lines was adjusted so that overlapping lines appear as a blend of the overlapping colours.

In the cases where metabolites A and B diffuse instantaneously, the CV is always zero as expected. For most other rate-limiting regimes, the CV does not generally change much with the PA ratio, and there are overlaps among regimes for some combinations of enzyme distribution. However, CV tends to be slightly sensitive to the PA ratio when both metabolites cannot diffuse. Specifically, CV increases at decreasing PA ratio. As previously discussed, growth only occurs in regions in the outermost layer when both metabolites cannot diffuse. At low PA ratios, only a small number of cell regions actually contribute to overall growth. The contribution of each region depends on its enzyme concentrations, some of which are sampled randomly. Since variance is usually high when sample size is small, the variance of regional growth among a small number of contributing regions tends to be high (Hajagos and Steiner, 2001). Consequently, cell growth which is the sum of these highly variable regional growth can take a wide range of values; thus CV tends to be higher at low PA ratios.

In the simulations so far, the reaction network and parameter values were designed such that no specific enzyme would be limiting growth if simulated as a cell-scale representation. In other words, there are just enough of each enzyme to support a specific cell growth level assuming that all reactions are not diffusion-limited. We called this the “default model”. When the “default model” is simulated as a regional-scale representation, some regions may have disproportional enzyme contents depending on how each enzyme is distributed. As diffusion becomes limiting, there are regions where some enzymes are in excess or unused. Consequently, the overall cell growth is lower than the theoretical maximum (Fig. 3). How much growth is reduced thus indicates the degree to which enzymes are not fully utilised. At the population level, a high CV indicates that a large proportion of cells under-utilise their enzymes and grow much lower than the maximum and mean levels. Therefore, the CV measures the extent of under-utilisation within the population.

We also explored alternative models where some enzymes were in excess even when diffusion is not limiting, by changing the stoichiometry in the reaction network while maintaining total cellular amount of each enzyme. We chose this over changing enzyme amount directly so that enzyme distribution patterns were consistent and comparable across models. In the first alternative which we called the “B-limiting model”, we increased the stoichiometric coefficient of metabolite B in reaction R4 as follows:

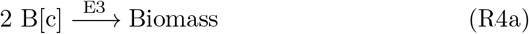

This was implemented by changing the element in the fourth column and third row in ***S*** in equation (3) to -2. Note that we are not indicating metabolite B is a growth-limiting nutrient. With this change, E3 becomes an excessive enzyme. Compared to the “default model”, there is not much difference in CV sensitivity to the PA ratio (Fig. 6). The most noticeable difference is a decrease in CV for several combinations of enzyme distribution in the regime where only metabolite A can diffuse instantaneously. We also tested a second alternative which we called the “A-limiting model”, where we increased the stoichiometric coefficient of metabolite A in reaction R3 instead (third column and second row of ***S***), as follows:

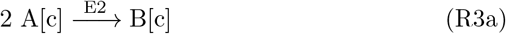

**Fig. 6.**
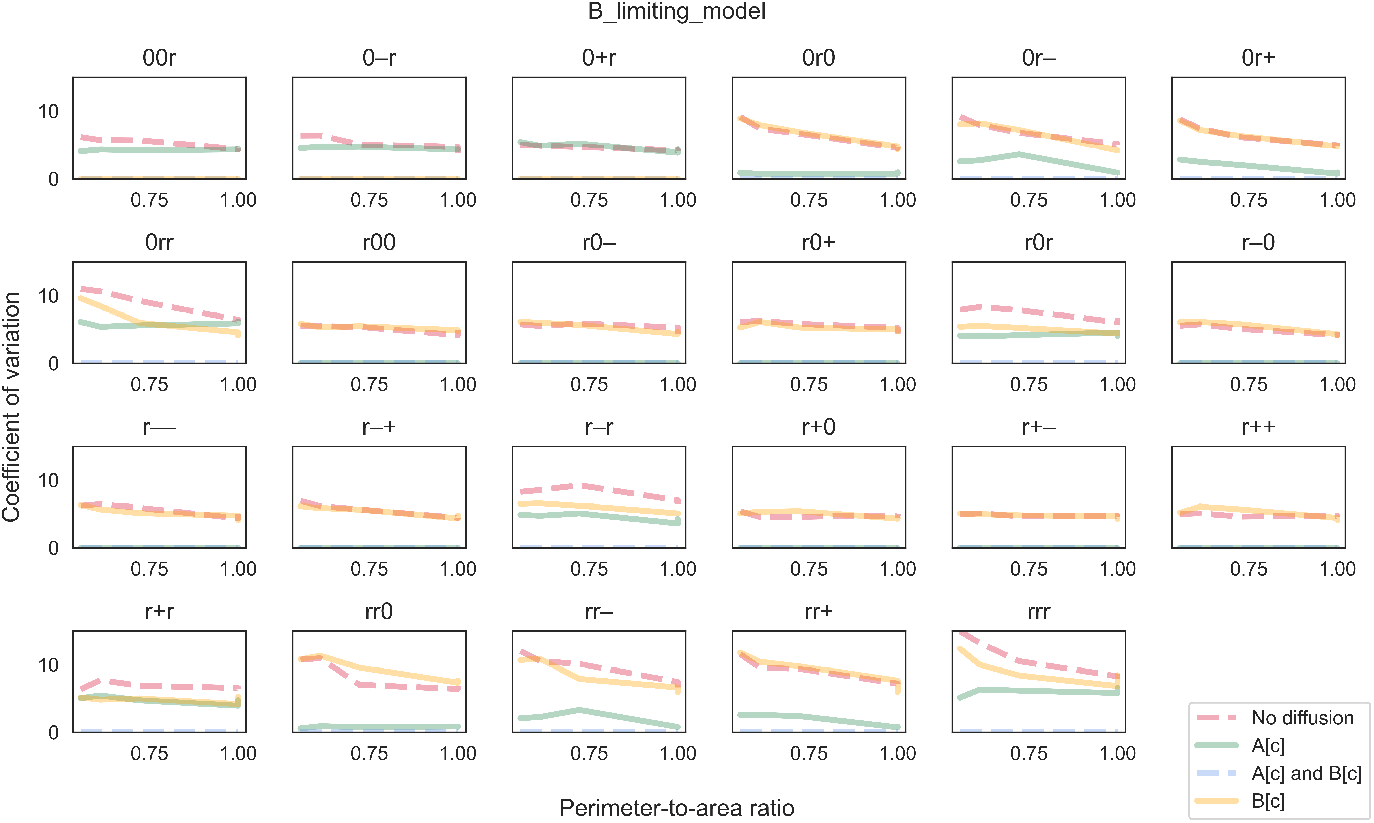
The coefficient of variation (CV) of biomass growth simulated using the “B-limiting model”. CV is plotted against perimeter-to-area ratio for different rate-limiting regimes (no diffusion, metabolite A can diffuse instantaneously, metabolite B can diffuse instantaneously, both metabolites can diffuse instantaneously). Each panel illustrates plots for a different combination of enzyme distribution patterns, as indicated by three characters at the top of the panel (“r”: random distribution; “0”: uniform distribution; “+”: positive gradient; “-”: negative gradient) that represent the distribution of transporter E1, enzyme E2 and enzyme E3, respectively. Opacity of the lines was adjusted so that overlapping lines appear as a blend of the overlapping colours.

In the “A-limiting model”, both enzymes E2 and E3 become excessive. CV becomes zero in all cases where metabolite A can diffuse instantaneously, regardless of whether B can diffuse (Fig. 7). More interestingly, the sensitivity of CV to the PA ratio increases for the other rate-limiting regimes, i.e. CV is high at low ratios and low at high ratios. In a number of cases, CV even approaches zero at the ratio of one, going from highly stochastic to almost deterministic just by changing the PA ratio. Together, our simulations suggest that enzyme distribution patterns within individual cells could affect not only single-cell behaviour but also variability at the population level. Besides ratelimiting regime and cell shape, effect at the population level also depends on the overall relative abundance of enzymes.

**Fig. 7.**
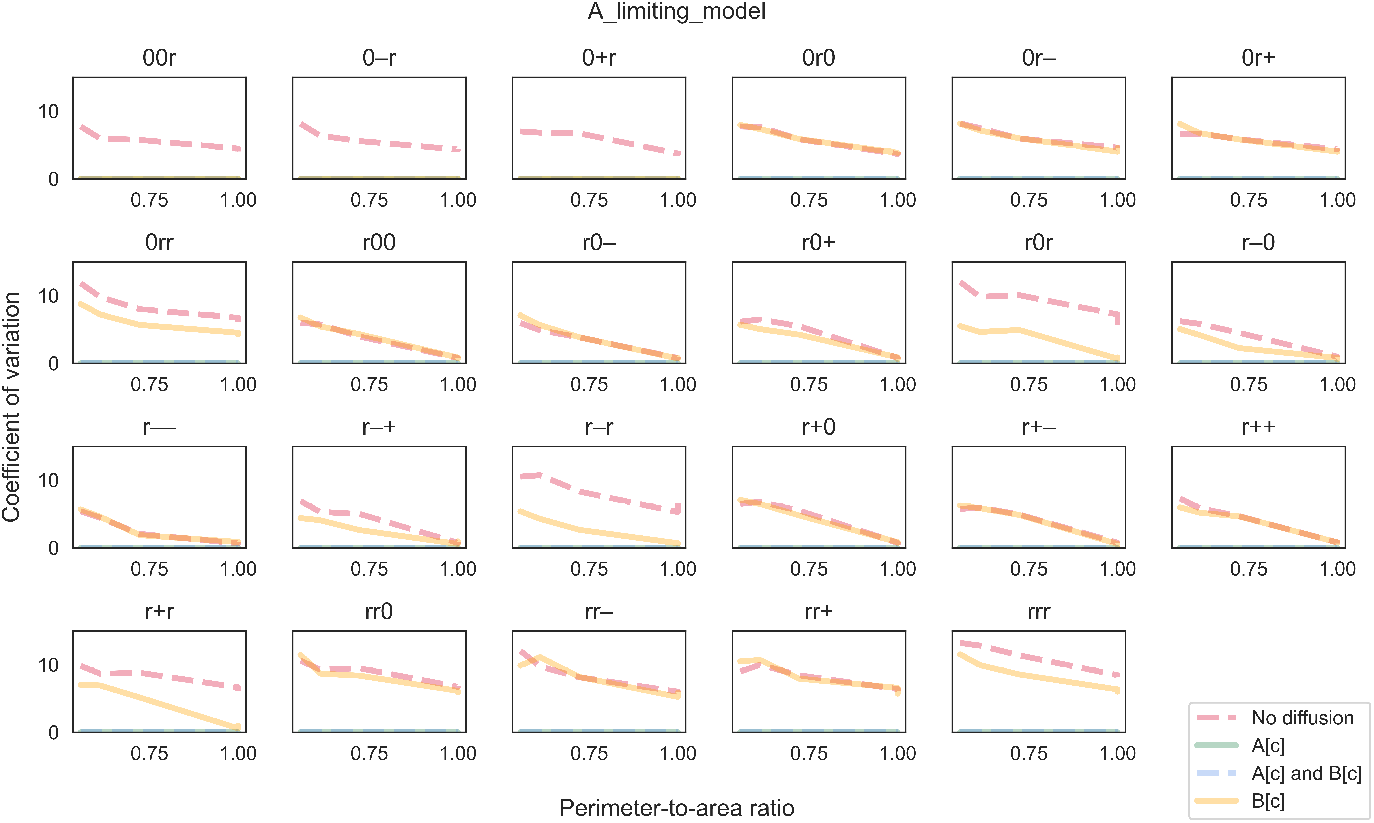
The coefficient of variation (CV) of biomass growth simulated using the “A-limiting model”. CV is plotted against perimeter-to-area ratio for different rate-limiting regimes (no diffusion, metabolite A can diffuse instantaneously, metabolite B can diffuse instantaneously, both metabolites can diffuse instantaneously). Each panel illustrates plots for a different combination of enzyme distribution patterns, as indicated by three characters at the top of the panel (“r”: random distribution; “0”: uniform distribution; “+”: positive gradient; “-”: negative gradient) that represent the distribution of transporter E1, enzyme E2 and enzyme E3, respectively. Opacity of the lines was adjusted so that overlapping lines appear as a blend of the overlapping colours.

### 5.4 Effect on robustness

Simulations of the linear-pathway example presented earlier have shown that cell shape and enzyme distribution pattern can affect the ability of cells to convert nutrients into final products such as biomass. These effects are due to diffusion and can be seen at both the individual and population levels. Using our method, we also investigated these effects in a branched-pathway example where nutrient can be converted into energy via an efficient primary path or a secondary path that also produces a toxic compound. Similar to the biomass objective in the first example, the maximised value of the objective function in this example is generally reduced at low PA ratios when either or both metabolites X and Y have limited ability to diffuse (Fig. 8). Again, the extent of reduction is strongest when both metabolites cannot diffuse, while the maximal value of 400 can be achieved regardless of the ratio when both metabolites can diffuse instantaneously. As for enzyme distribution, the effect of diffusion is only influenced by the distribution patterns of transporter E4, enzyme E5 and enzyme E6. The spatial distribution of enzyme E7 that catalyses the secondary path has negligible influence. This is not surprising given that the secondary path is not favourable and rarely used (Fig. 10). Besides analysing the maximised value of the objective function that includes a penalty for toxin production, we also examined the effect of diffusion on energy production alone (Fig. 9). By comparing the slope of lines in Fig. 8 and Fig. 9, it can be seen that the total amount of energy produced is less sensitive to PA ratio. In fact, when metabolite X can diffuse instantaneously, PA ratio has almost no effect on total energy production even when the objective value is sensitive. Simulated cells achieve this by diverting some portion of metabolite Y to utilising the secondary path for energy production (Fig. 10). A similar scenario can also be observed when both metabolites X and Y have limited ability to diffuse, though this only occurs for some combinations of enzyme distribution patterns and the decrease in sensitivity is not as large. Thus, utilising the secondary path can help buffer against diffusion limitation. However, this comes at the cost of toxin production so the trade-off is only advantageous under certain conditions. Note that the extent of trade-off and the conditions under which it is advantageous depend on the objective function, i.e. the choice of objective coefficient or weight that determines the amount of penalty incurred.

**Fig. 8.**
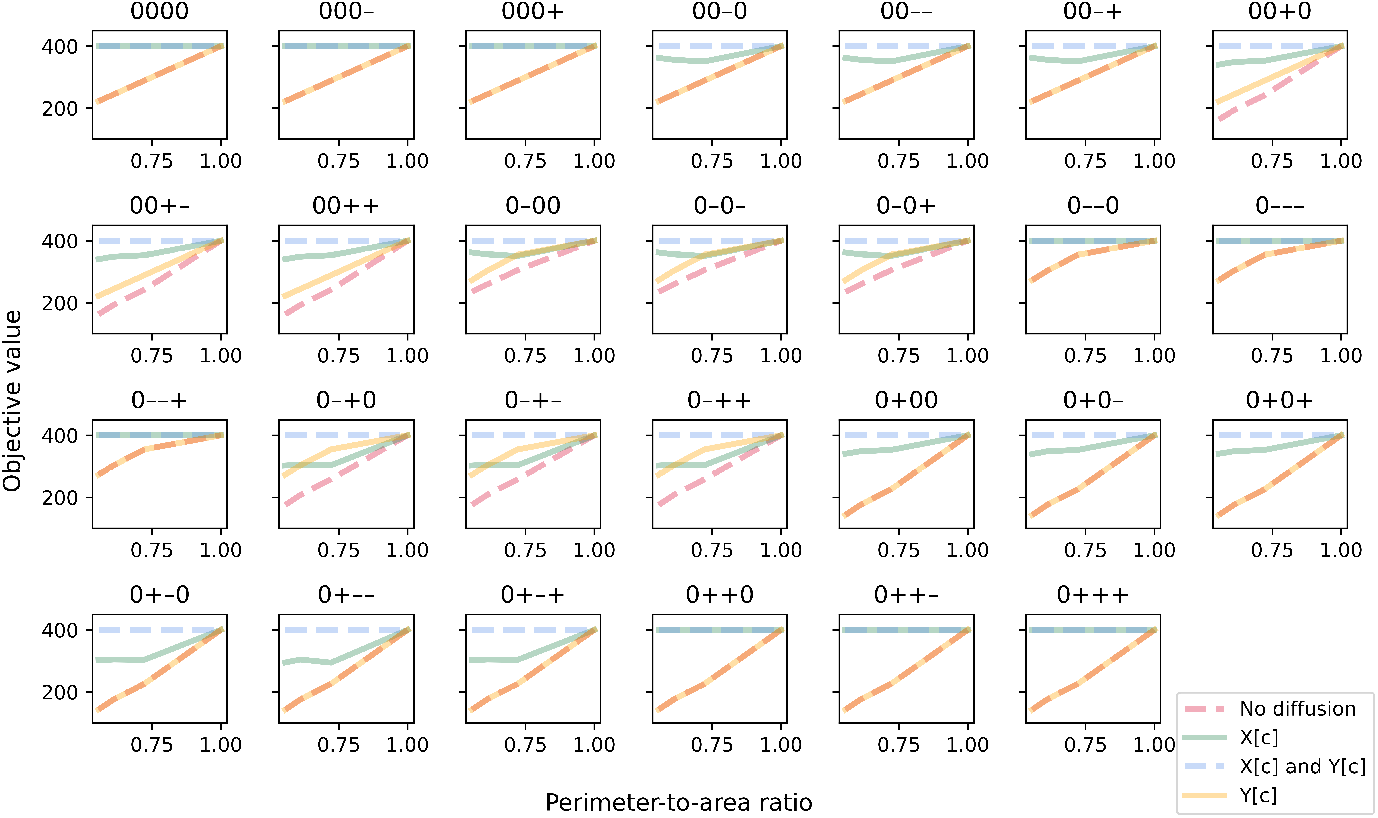
Maximised value of the objective function in the branched-pathway model, plotted against perimeter-to-area ratio for different rate-limiting regimes (no diffusion, metabolite X can diffuse instantaneously, metabolite Y can diffuse instantaneously, both metabolites can diffuse instantaneously). Each panel illustrates plots for a different combination of enzyme distribution patterns, as indicated by four characters at the top of the panel (“0”: uniform distribution; “+”: positive gradient; “-”: negative gradient) that represent the distribution of transporter E4, enzyme E5, enzyme E6 and enzyme E7, respectively. Opacity of the lines was adjusted so that overlapping lines appear as a blend of the overlapping colours.

**Fig. 9.**
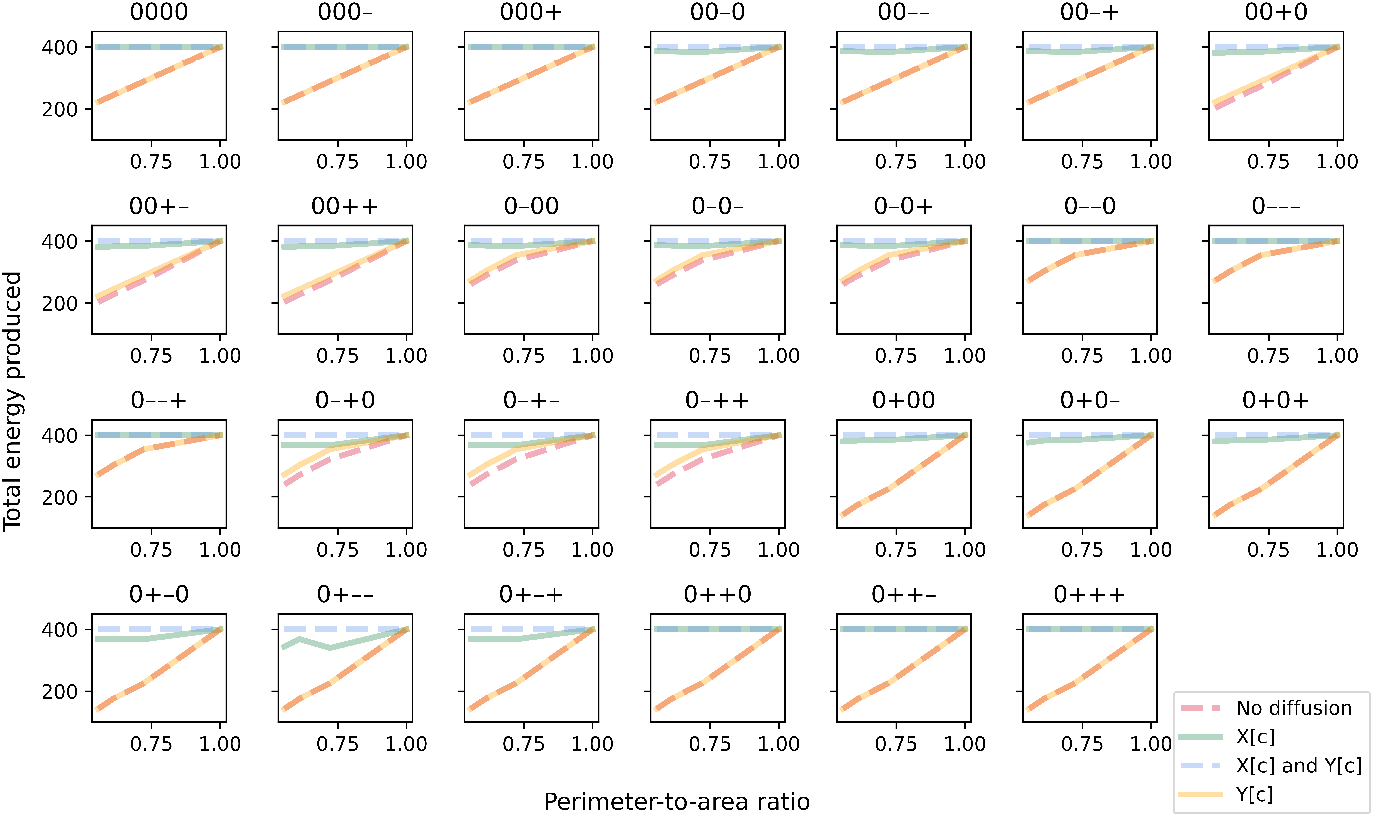
Simulated energy production plotted against perimeter-to-area ratio for different rate-limiting regimes (no diffusion, metabolite X can diffuse instantaneously, metabolite Y can diffuse instantaneously, both metabolites can diffuse instantaneously). Each panel illustrates plots for a different combination of enzyme distribution patterns, as indicated by four characters at the top of the panel (“0”: uniform distribution; “+”: positive gradient; “-“: negative gradient) that represent the distribution of transporter E4, enzyme E5, enzyme E6 and enzyme E7, respectively. Opacity of the lines was adjusted so that overlapping lines appear as a blend of the overlapping colours.

**Fig. 10.**
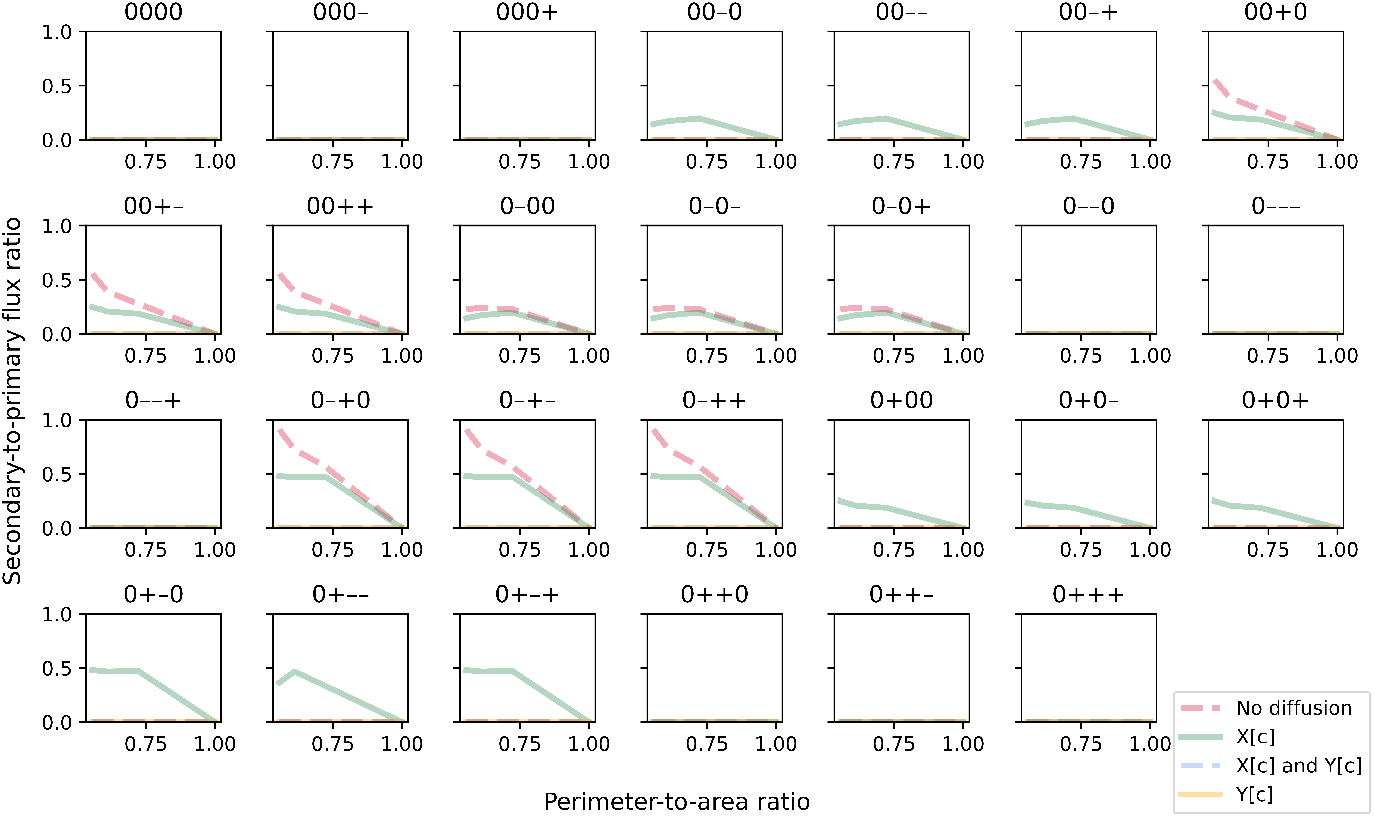
The secondary-to-primary flux ratio plotted against perimeter-to-area ratio for different rate-limiting regimes (no diffusion, metabolite X can diffuse instantaneously, metabolite Y can diffuse instantaneously, both metabolites can diffuse instantaneously). Each panel illustrates plots for a different combination of enzyme distribution patterns, as indicated by four characters at the top of the panel (“0”: uniform distribution; “+”: positive gradient; “-”: negative gradient) that represent the distribution of transporter E4, enzyme E5, enzyme E6 and enzyme E7, respectively. Opacity of the lines was adjusted so that overlapping lines appear as a blend of the overlapping colours.

We also found that for each rate-limiting regime, combinations of enzyme distribution can be grouped based on their effective maximal energy production (Fig. 11). For the regime where only metabolite X can diffuse instantaneously (left panel in Fig. 11), a maximal energy production of 400 can still be achieved solely through the primary path, i.e. zero secondary-to-primary flux ratio, as long as enzymes E5 and E6 have the same distribution pattern. The reason for this is similar to the linear-pathway example. Since X can move freely and instantaneously, it can theoretically occupy each cell region at a rate that enables enzyme E5 in every region to operate at maximal capacity, to convert X to Y without being limited by diffusion. When both enzymes have the same distribution pattern, there is proportional amount of regional enzyme E6 to convert Y to energy on site, so the limited ability of Y to diffuse has no effect on energy production. Interestingly, this diffusion limitation of Y has very little effect even in other combinations of enzyme distribution pattern. The largest decrease in energy production for this diffusion regime is less than 15%, with concomitant increase in the secondary-to-primary flux ratio. This reiterates the advantage of partial diversion to an alternative path. However, this path is not utilised at all if Y can diffuse instantaneously while X has limited ability to diffuse (middle panel in Fig. 11). This means that, at the level of penalty that we used in the simulation, toxin production is too high a cost to pay in this diffusion regime under any combinations of enzyme distribution pattern. On the other hand, if both metabolites have limited ability to diffuse (right panel in Fig. 11), three groups of enzyme distribution patterns can benefit from partial diversion to the alternative path but not the other three. The largest secondary-to-primary flux ratio is seen in this diffusion regime, going up to 0.9 in the case where enzyme E5 is spatially distributed on a positive gradient while enzyme E6 is on a negative gradient. This flux ratio enables simulated cells to generate about 88% of the maximal energy production achievable in this diffusion regime.

**Fig. 11.**
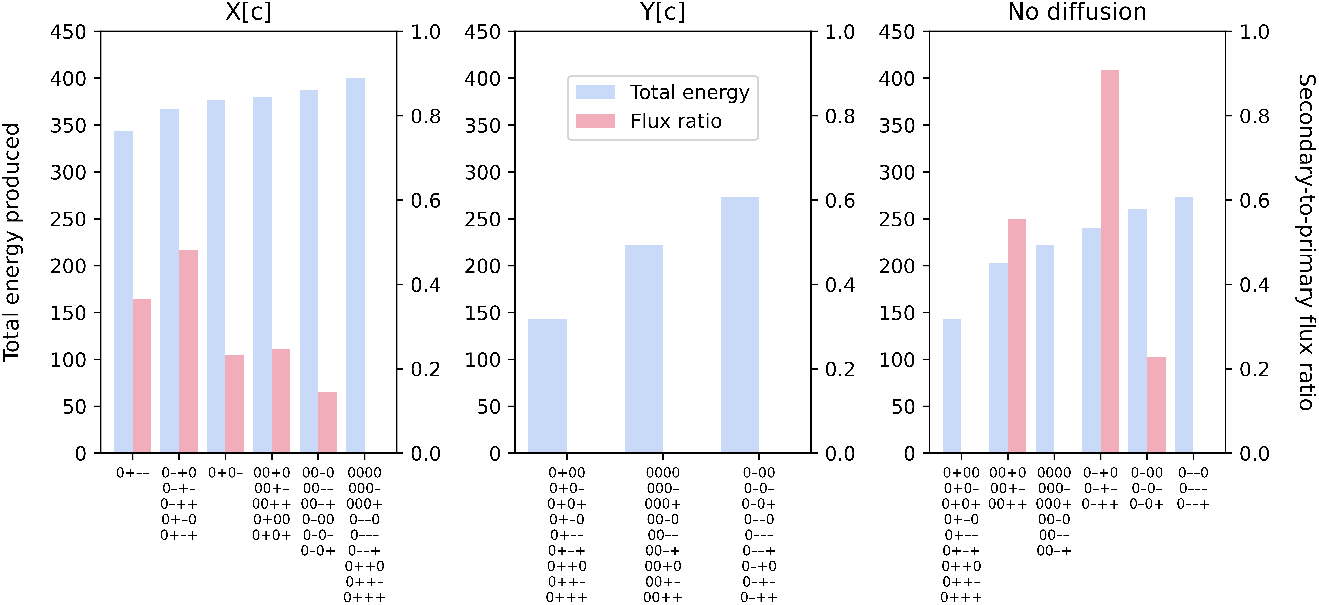
Simulated energy production (blue bars, left axis) and secondary-to-primary flux ratio (red bars, right axis) for a 6x6 cell for different rate-limiting regimes: only metabolite X can diffuse instantaneously (left panel); only metabolite Y can diffuse instantaneously (middle panel); and both metabolites are restricted (right panel). The combinations of enzyme distribution patterns that produce each growth level are indicated at the bottom of each bar. Each combination of four characters (“0”: uniform distribution; “+”: positive gradient; “-”: negative gradient) represents the distribution of transporter E4, enzyme E5, enzyme E6 and enzyme E7, respectively.

For stochastic simulations that involve at least one enzyme that is spatially distributed in a random pattern, the CV for both objective value (Fig. 12) and total energy produced (Fig. 13) are always zero in the cases where metabolites X and Y can diffuse instantaneously, as expected. Similar to the linear-pathway example, CV tends to be slightly sensitive to the PA ratio when both metabolites cannot diffuse, with CV increasing at decreasing PA ratio. On the other hand, when only metabolite Y has a limited ability to diffuse, this sensitivity is lost for both objective value and total energy produced. We also observed that the CV for total energy produced is generally lower than that for objective value and approaches zero in many cases. Mathematically, the CV can differ between the two only if the secondary path is utilised. This means that the observed reduction in both cell-to-cell variability of total energy produced and the sensitivity of this variability to cell shape is due to partial diversion to the alternative path. Altogether, our results show that a branched pathway, where there is an alternative path to generating the same desirable products, can provide robustness to the system. This concurs with a previous study demonstrating that enhanced robustness in metabolic networks is related to the organisation of branched metabolites (Smart et al, 2008). Our results further suggest that factors such as cell shape, diffusion regime and spatial distribution of enzyme can influence this robustness, at both single-cell and population levels.

**Fig. 12.**
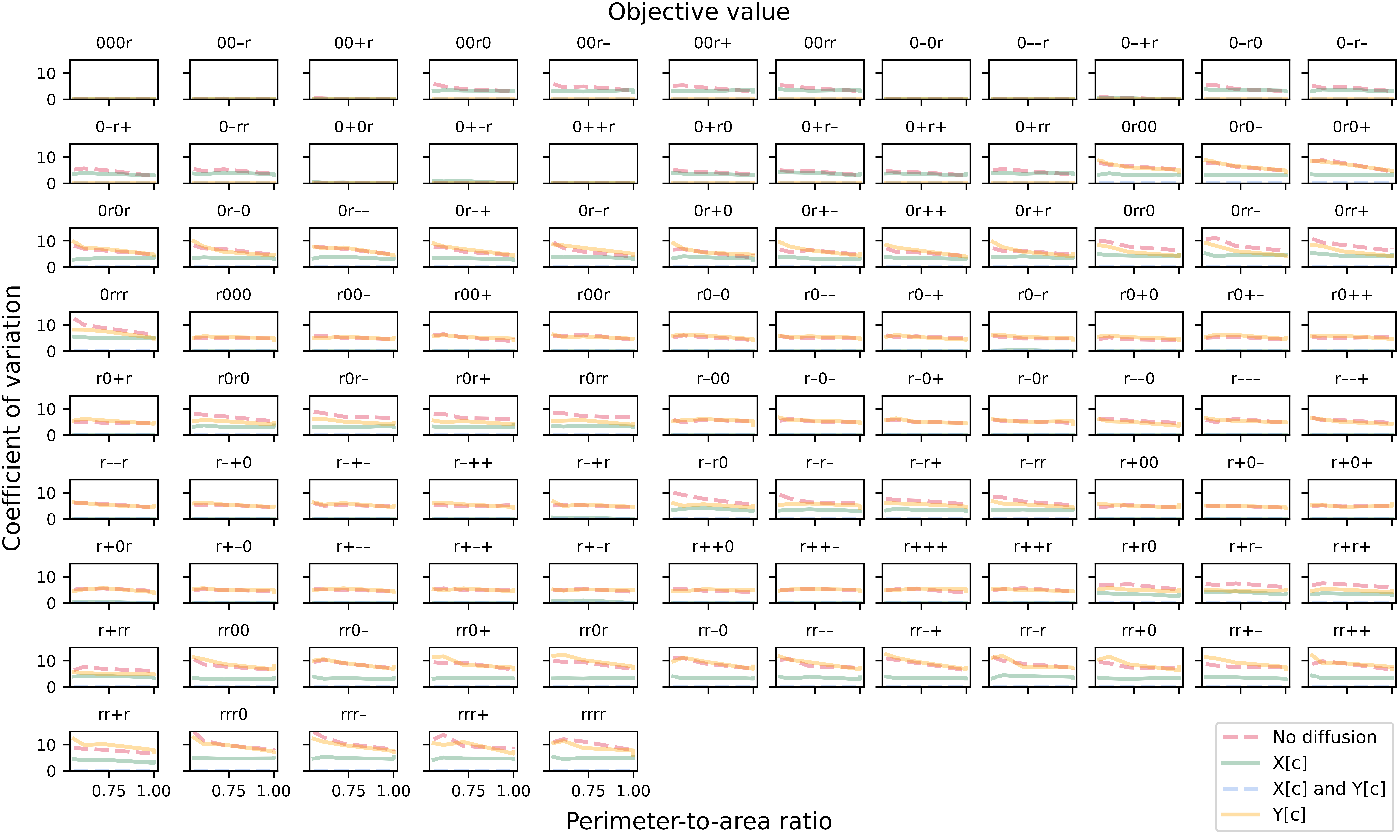
The coefficient of variation (CV) of maximised objective value in the branched-pathway model. CV is plotted against perimeter-to-area ratio for different rate-limiting regimes (no diffusion, metabolite X can diffuse instantaneously, metabolite Y can diffuse instantaneously, both metabolites can diffuse instantaneously). Each panel illustrates plots for a different combination of enzyme distribution patterns, as indicated by four characters at the top of the panel (“r”: random distribution; “0”: uniform distribution; “+”: positive gradient; “-”: negative gradient) that represent the distribution of transporter E4, enzyme E5, enzyme E6 and enzyme E7, respectively. Opacity of the lines was adjusted so that overlapping lines appear as a blend of the overlapping colours.

**Fig. 13.**
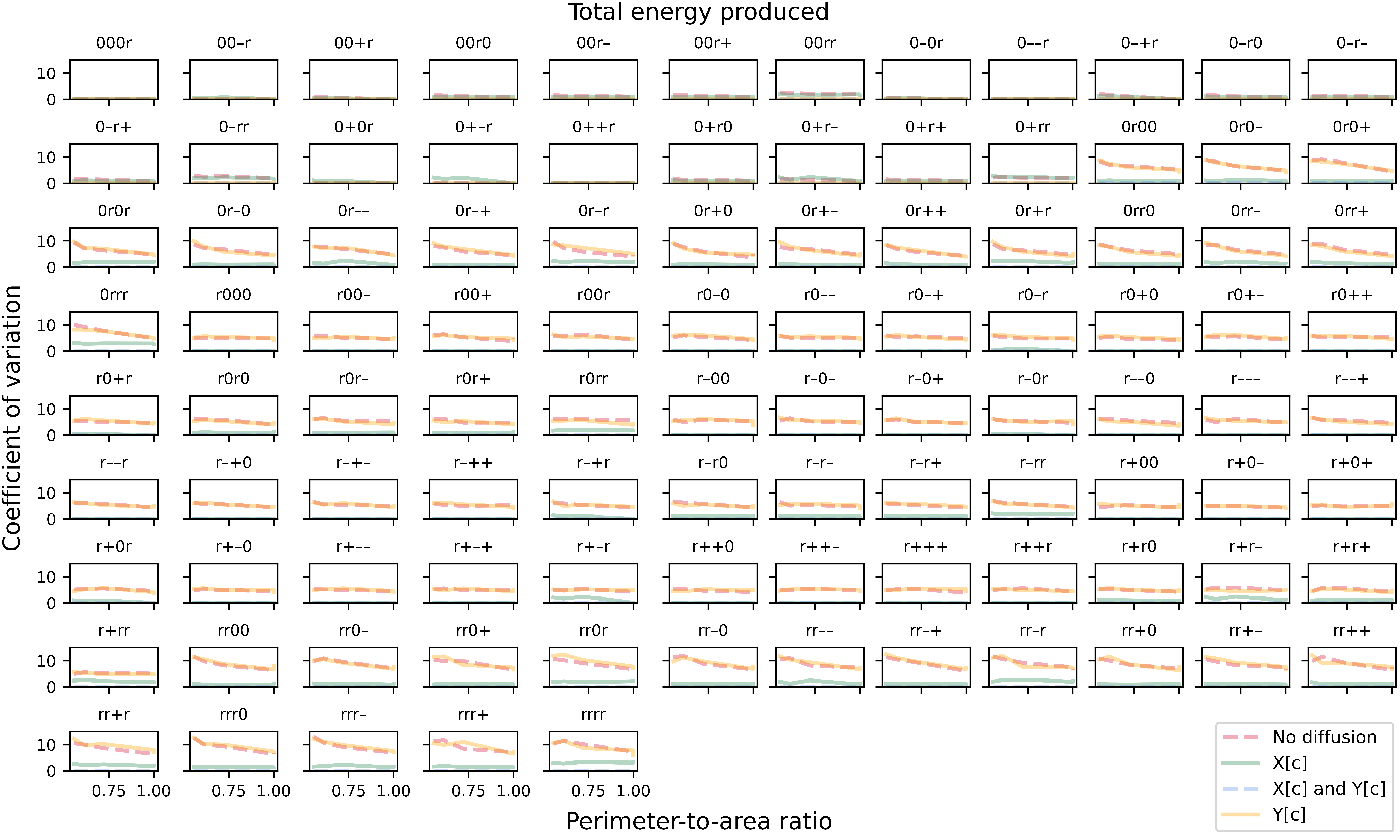
The coefficient of variation (CV) of total energy produced. CV is plotted against perimeter-to-area ratio for different rate-limiting regimes (no diffusion, metabolite X can diffuse instantaneously, metabolite Y can diffuse instantaneously, both metabolites can diffuse instantaneously). Each panel illustrates plots for a different combination of enzyme distribution patterns, as indicated by four characters at the top of the panel (“r”: random distribution; “0”: uniform distribution; “+”: positive gradient; “-”: negative gradient) that represent the distribution of transporter E4, enzyme E5, enzyme E6 and enzyme E7, respectively. Opacity of the lines was adjusted so that overlapping lines appear as a blend of the overlapping colours.

## 6 Discussion

In this paper, we present discretised FBA, a spatial modelling method and algorithms that extend conventional FBA to include representation of metabolite diffusion inside cells. Implemented on a grid-based system, our method allows us to spatially explore metabolism and cell behaviour such as growth when diffusion becomes a limiting factor in enzymatic reactions. Coupling FBA with spatial representation has been done previously, for example to describe the interactions and spatial dynamics among microbial communities (Henson, 2015; Borer et al, 2019; Karimian and Motamedian, 2020; Dukovski et al, 2021). To the best of our knowledge, our work is the first spatially informed FBA method that looks into reaction-diffusion within a single cell. Compared to existing spatial modelling methods such as PDE, our method requires very few parameters.

We simulated diffusion-limited and reaction-limited regimes using extreme settings where metabolites are either restricted or able to diffuse freely and instantaneously. This allowed us to compare the maximum extent of their potential impact on behaviour such as growth and energy production. In real systems, metabolites are generally small so they can move around quite freely and rapidly within the same compartment, but can be impeded by surrounding macromolecules. Even so, it is unlikely that metabolites will be fully restricted to the extreme level of diffusion limitation in our simulations. Besides, highly efficient enzymes that are diffusion-limited are rare (Sweetlove and Fernie, 2018). The majority of enzymes have not evolved to the “perfect” level of catalytic efficiency, so most reactions likely operate under a reaction-limited regime even when the diffusion rates of substrates are low (Bar-Even et al, 2015). Additionally, metabolism consists of reaction networks that are much more complex than our examples, so we expect the overall impact is intricate but much less severe. Some studies have proposed that diffusion limitation can be overcome through metabolite channelling, where intermediate products in a series of reactions are directly transferred from one enzyme to the next in the pathway to shorten diffusion distance (Obata, 2020). All these seem to indicate a negligible effect of metabolite diffusion on single-cell metabolism. However, several studies have noticed that the density of some bacterial cells seem to be optimal for achieving cellular efficiency, with a trade-off between high volume density that can improve biochemical efficiency and molecular crowding that can slow down diffusion (Pang and Lercher, 2023; Vazquez, 2010). Since the sub-cellular environment is not static, spatial and structural organisation in cells could provide another dimension of regulation. Our method provides a theoretical means to study the relationship between local reaction/diffusion limitations and optimal behaviour such as growth, energy production or any other objective function. This relationship could be useful, for example to study cancer cells that are known to have abnormal growth and display drug resistance predictable by cell shape (Longden et al, 2021). Another potential use is to facilitate the design of synthetic systems, such as protocells where structure is a major consideration (Gözen et al, 2022).

We also demonstrated another utility of our spatial method by presenting simulations of different enzyme distribution patterns. Experimental observations have shown that most cellular molecules are heterogeneously distributed. Such heterogeneity could arise as the direct effect of regulation and the indirect effect of cell shape (Halatek et al, 2018; Ovádi and Saks, 2004). Since enzymes are usually large and metabolism is a relatively fast process, we assumed that enzymes do not move during the time-scale of our steady-state simulations. Our results suggest that some distribution patterns are more favourable for growth and energy production than others. For linear pathways, the more favourable distributions are those where enzymes that act in series have similar range of concentration or capacity within the same region. This supports the notion that cells have organelles and compartments, among other theories, to concentrate enzymes and substrates that participate in the same pathways to increase biochemical efficiency (Diekmann and Pereira-Leal, 2013; Conrado et al, 2007). For branched pathways, some combinations of enzyme distribution patterns and diffusion regimes favour a partial diversion of branch-point metabolites to the alternative paths, even when there is a large incurred cost. In our example, this cost comes in the form of toxin production that could be detrimental to cell growth and survival. The detrimental effect of toxin can occur directly and indirectly through metabolism as well as other cellular processes. We demonstrated one way of representing this cost, through the use of multi-objective optimisation. This can be further explored in future, for example by varying the weight of each objective, which could be useful for helping design cancer therapy (Lee et al, 2020). A recent work made promising progress in engineering cells with specific patterns of protein localisation, so it may soon be possible to experimentally test theoretical effects of enzyme distribution patterns (Rajasekaran et al, 2024).

Our results also suggest that enzyme distribution within individual cells can affect variability at the population level, if some reactions are diffusion-limited. Enzymes are required for cell growth but their production is costly (Buttgereit and Brand, 1995; Russell and Cook, 1995). In our simulations, the same investment, i.e. the same amount of total cellular enzyme, could yield different returns in growth and energy production if cells change their shape and enzyme distribution pattern. At the population level, a low CV indicates a trade-off between overall return (population growth and energy production) and low cell-to-cell variability. On the other hand, a high CV means that a small number of cells in the population yield unusually high (or low) growth and energy, which could confer a selective advantage under fluctuating environments (Balaban et al, 2004; Acar et al, 2008). Most studies on variability have focused on stochasticity in gene expression (Thattai and van Oudenaarden, 2004; Schwabe et al, 2011; Evans and Zhang, 2020); our study suggests that other aspects such as cell shape and spatial organisation may have effects under certain conditions.

## 7 Conclusions and future work

In conclusion, our method expands the widely used FBA and increases its spatial resolution. FBA is widely used because it has low parameter requirements yet provides flexibility for investigation. For example, one can use thermodynamics principles and gene expression data to constrain reaction fluxes and direction when kinetic parameters are lacking at the genome scale, or test various hypotheses by changing the objective function (Blazier and Papin, 2012; Machado and Herrgard, 2014; Raman and Chandra, 2009). We recommend similar use of our method. In the absence of data about diffusion coefficients and concentrations, spatial information and measurements of factors that affect diffusivity can be used to constrain diffusion fluxes, to gain insight into the system.

Currently, our method is implemented to represent regular two-dimensional shapes. We have demonstrated the use of this method on two simple toy reaction networks in single-compartment cells. Future extensions could include capabilities to represent more complex cell shapes and organelles, as well as in three dimensions. At present, each cell region or groups of neighbouring regions can effectively be a distinct sub-cellular compartment or organelle, but all movements of metabolites in and out of a region are treated as diffusion. To represent membrane-bound organelles, extra features are needed to identify cell regions located at the boundary of an organelle, and treat metabolite movements between these regions and neighbouring regions that are not part of the organelle as cross-membrane (active/passive) transport reactions. Additional functions could facilitate the representation of organelle-specific reaction networks. These features will enable us to simulate more realistic cells and reaction networks, so that we can explore how spatiotemporal organisation of sub-cellular structure affects metabolism and cell behaviour such as growth.

## Acknowledgments

We thank Alexandra-Anamaria Sorinca who tested the idea behind this method as a summer undergraduate project funded by the London Mathematical Society (URB-2022-19). This work was funded by the United Kingdom Research Innovation Future Leaders Fellowship (MR/T043571/1).

## Appendix A

The following details how our Python API converts a cell-scale representation of metabolic network into a regional-scale representation (see Algorithms 1–5):

1. Create a two-dimensional *I* x *J* grid, where *I* (cell width) and *J* (cell length) are defined by the user;
2. Create in every cell region a metabolite object for each intracellular metabolite. For metabolites in the extracellular compartment, only one object per metabolite will be created. The user has to provide input of metabolites as two separate lists; one for intracellular and another for extracellular metabolites;
3. Create in every cell region a reaction object for each intracellular biochemical reaction. For reactions in the extracellular compartment including exchange reactions, only one object per reaction will be created. Again, the user has to provide the two groups of reactions separately;
4. Create an objective function that sums over all cell regions, the variable(s) which represents the flux(es) of a user-specified intracellular reaction(s), e.g. the biomass reaction;
5. Create in every boundary cell region a reaction object that represents the transport of each intracellular metabolite to/from the extracellular compartment. Boundary cell regions are the regions located in the outermost layer of the cell, i.e. the boundary between cell and the extracellular compartment. For this step, the user has to provide the list of intracellular metabolites that are transported to/from the extracellular compartment, and the list of import/export reactions;
6. For each cell region:
  a. identify its neighbouring regions;
  b. for each neighbouring region, create a reaction object for every intracellular metabolite to represent diffusion from own region to the neighbouring region;
7. For each reaction object, set the bounds of its flux variable based on user input. If no bounds are specified for a reaction, its flux variable will be left unbounded.

Once the above metabolite and reaction objects have been created and their properties defined, the COBRApy package will automatically formulate the corresponding linear programming problem.

### Algorithm 1

Create cell regions

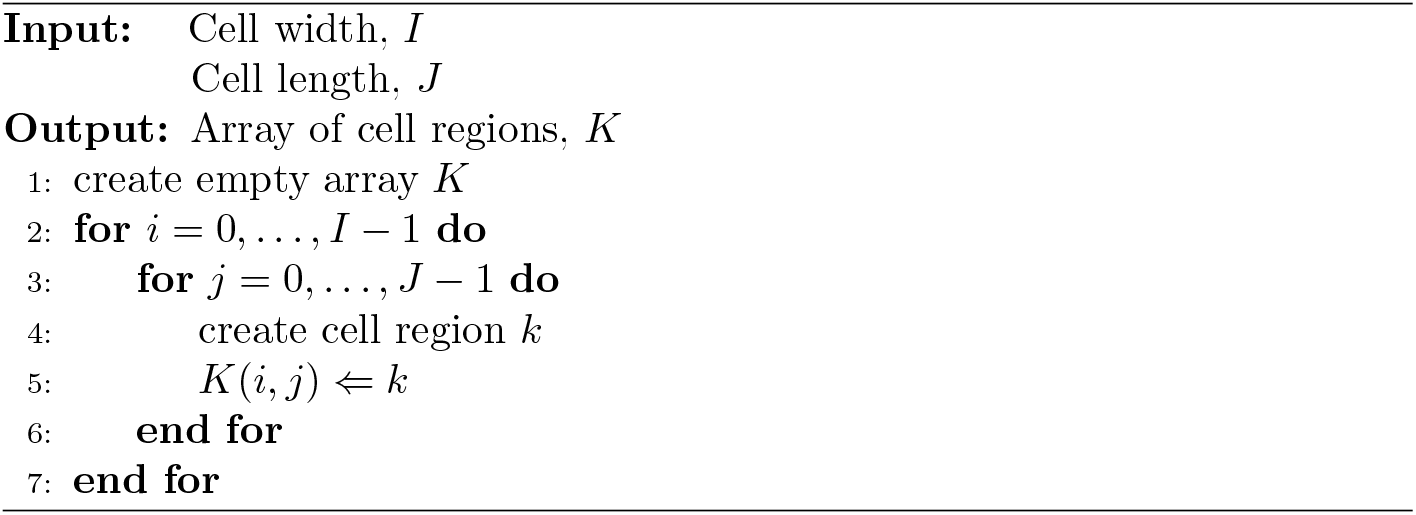

### Algorithm 2

Create regional metabolites

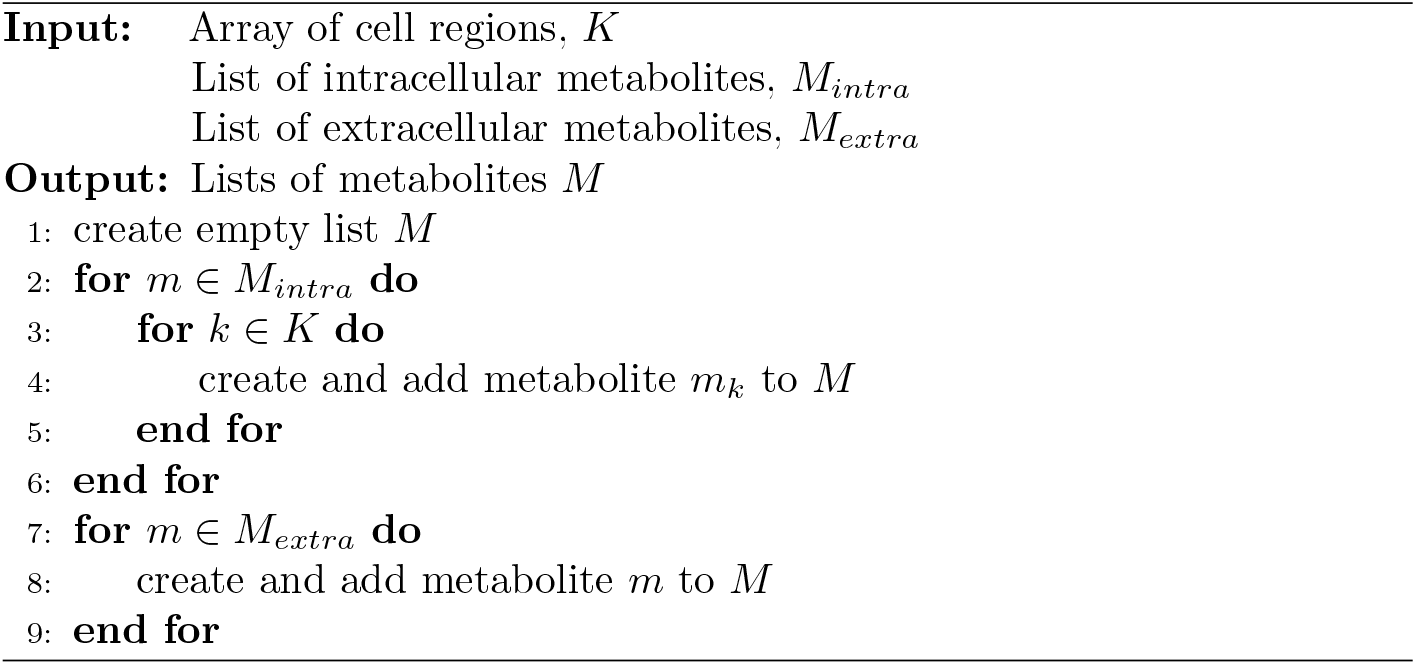

### Algorithm 3

Create regional reactions

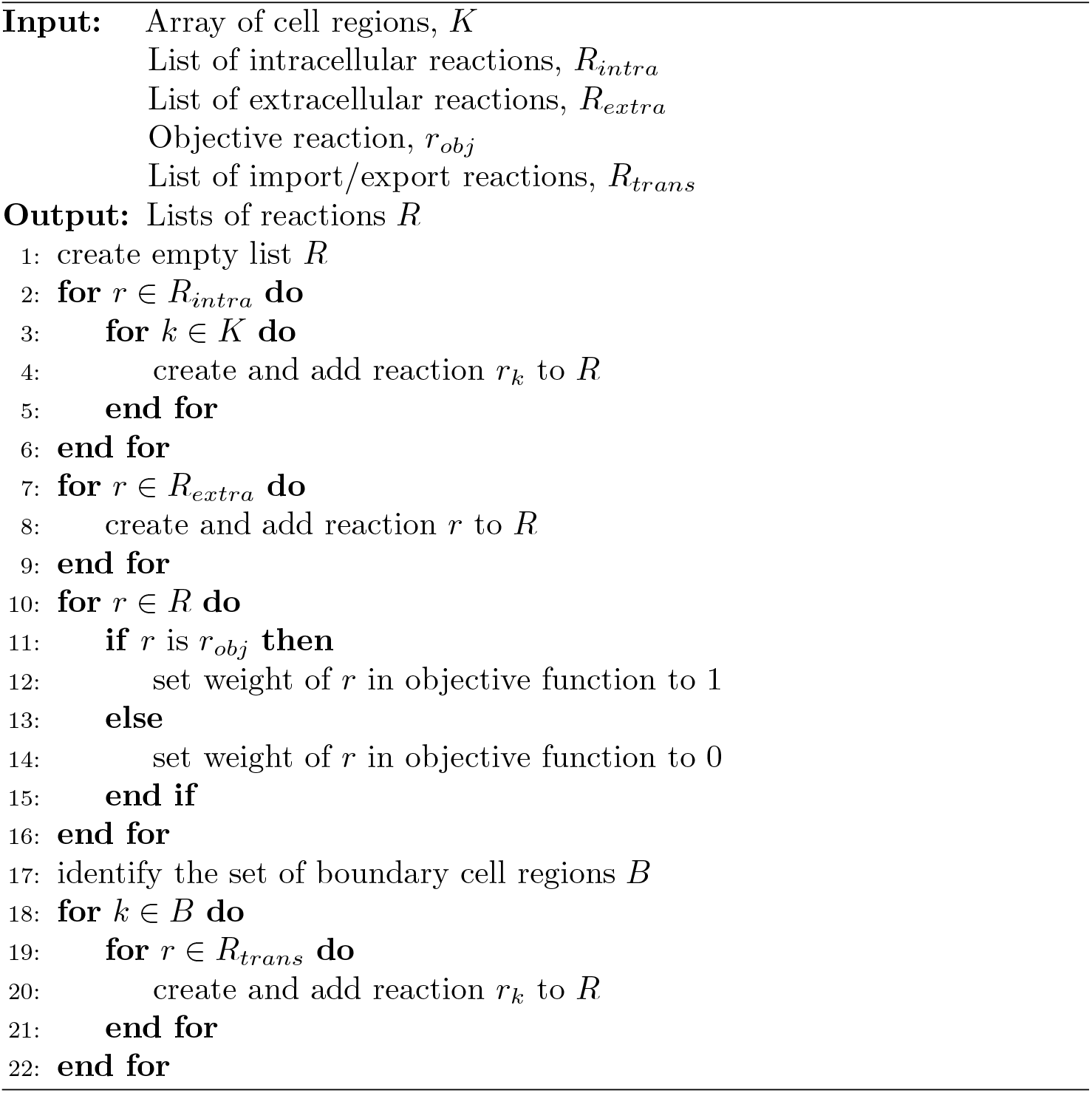

### Algorithm 4

Add metabolite diffusion to list of reactions

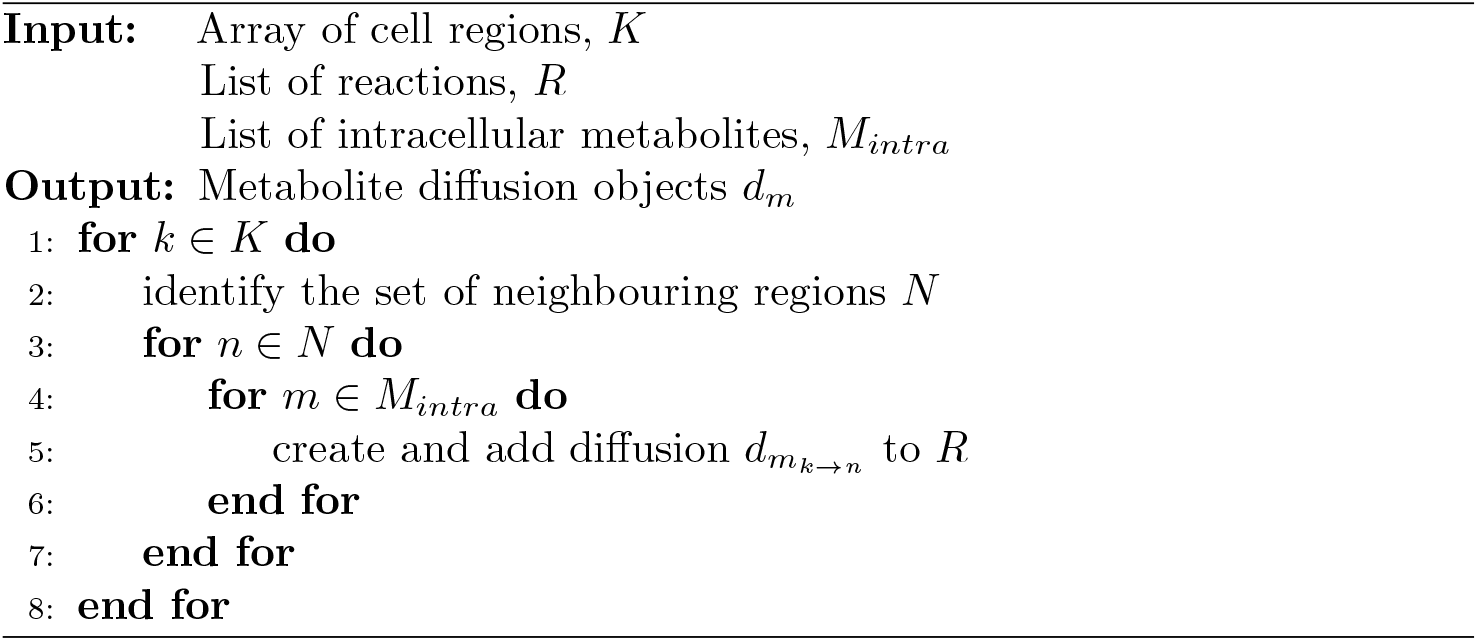

### Algorithm 5

Set reaction bounds

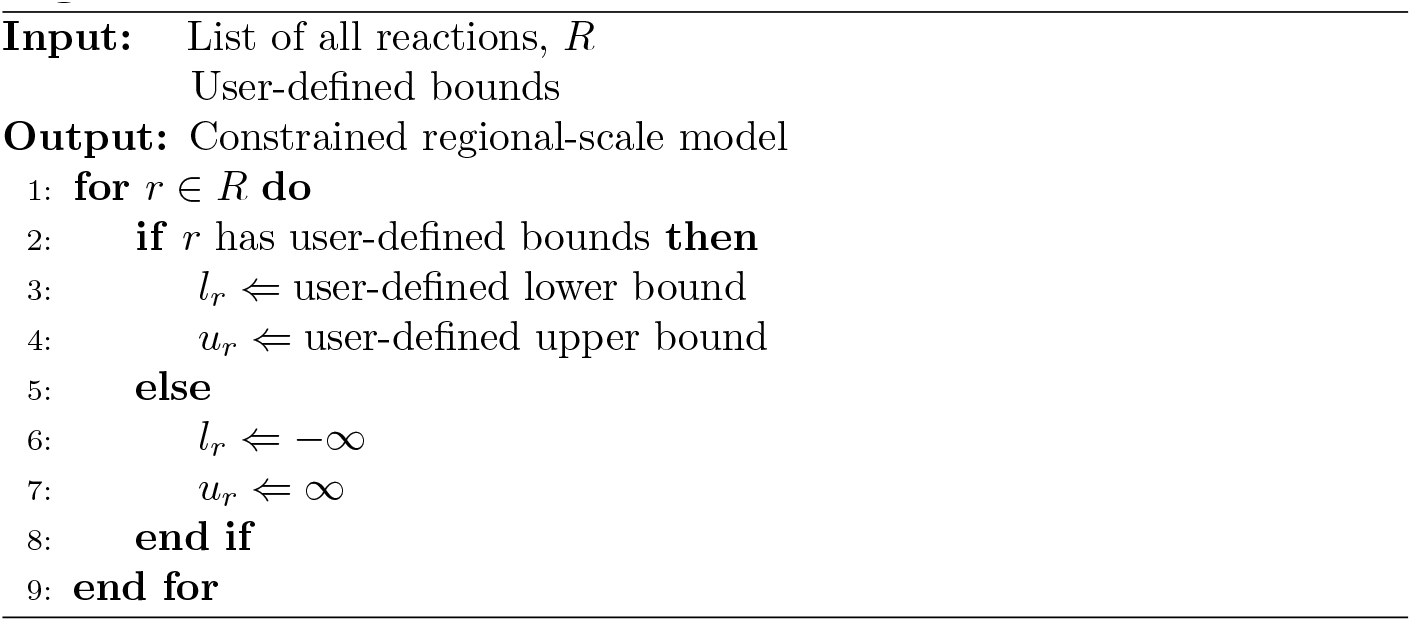

## Appendix B

To simulate different enzyme distribution patterns, we provide additional functions in the API that do the following:

1. Calculate the concentration of a user-specified enzyme in each cell region based on the cellular concentration and distribution pattern provided by users. The function provides the option of distributing enzymes in the cell uniformly, on a positive gradient (increasing concentration from the boundary to the centre of the cell), on a negative gradient, or randomly. Expected user inputs to determine the distribution pattern include whether the distribution is random, if distributing to all cell regions or only the outermost layer, the gradient parameter and an optional seed for the random number generator (see Algorithm 6);
2. Calculate the upper bound of each regional flux variable for the reaction catalysed by the enzyme, based on the regional concentration calculated above and a user-specified *k*_*cat*_ value.

### Algorithm 6

Distribute enzyme

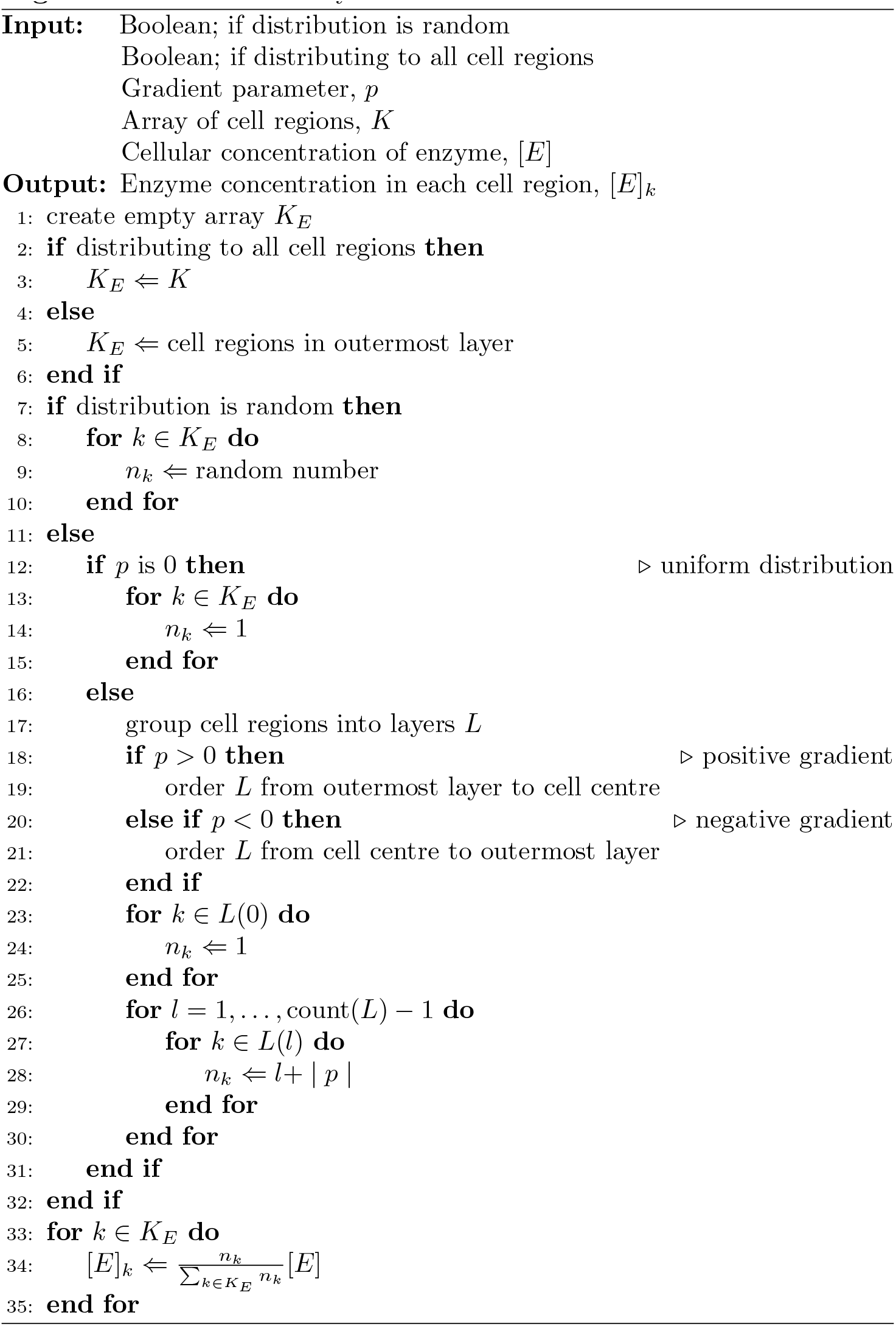

